# The dynamic, combinatorial cis-regulatory lexicon of epidermal differentiation

**DOI:** 10.1101/2020.10.16.342857

**Authors:** Daniel Kim, Viviana Risca, David Reynolds, James Chappell, Adam Rubin, Namyoung Jung, Laura Donohue, Arwa Kathiria, Minyi Shi, Zhixin Zhao, Harsh Deep, Howard Y. Chang, Michael P. Snyder, William J. Greenleaf, Anshul Kundaje, Paul A. Khavari

## Abstract

Transcription factors (TFs) bind DNA sequence motif vocabularies in cis-regulatory elements (CREs) to modulate chromatin state and gene expression during cell state transitions. A quantitative understanding of how motif lexicons influence dynamic regulatory activity has been elusive due to the combinatorial nature of the cis-regulatory code. To address this, we undertook multi-omic data profiling of chromatin and expression dynamics across epidermal differentiation to identify 40,103 dynamic CREs associated with 3,609 dynamically expressed genes, then applied an interpretable deep learning framework to model the cis-regulatory logic of chromatin accessibility. This identified cooperative DNA sequence rules in dynamic CREs regulating synchronous gene modules with diverse roles in skin differentiation. Massively parallel reporter analysis validated temporal dynamics and cooperative cis-regulatory logic. Variants linked to human polygenic skin disease were enriched in these time-dependent combinatorial motif rules. This integrative approach reveals the combinatorial cis-regulatory lexicon of epidermal differentiation and represents a general framework for deciphering the organizational principles of the cis-regulatory code in dynamic gene regulation.

**HIGHLIGHTS:**

- An integrative multi-omic resource profiling chromatin and expression dynamics across keratinocyte differentiation
- Predictive deep learning models of chromatin dynamics reveal a high-resolution cis-regulatory DNA motif lexicon of epidermal differentiation
- Model interpretation enables discovery of combinatorial cis-regulatory logic of homotypic and heterotypic motif combinations
- Massively parallel reporter experiments validate temporal dynamics and cis-regulatory logic of the combinatorial motif lexicon

## INTRODUCTION

The outermost layer of the skin, the epidermis, is formed and maintained by a dynamic homeostatic process involving the conversion of metabolically active basal cells that adhere to the epithelial basement membrane into cells that undergo cell cycle arrest and migrate outwards, engaging a program of terminal differentiation to form cornified keratinocytes (**Fig. S1A**). A host of human diseases are caused by disruption of this differentiation process (Lopez-Pajares et al., 2013). Calcium-induced differentiation of primary human keratinocytes *in vitro* mimics key properties of *in vivo* epidermal differentiation, making it a simple, tractable, and accurate *in vitro* system to study this medically relevant cellular differentiation.

Such differentiation processes involve dynamic cell state transitions accompanied by genome-wide changes in gene expression, chromatin state and three dimensional genome organization (Levine, 2010; Spitz and Furlong, 2012). Transcription factors (TFs) orchestrate these chromatin and expression dynamics by cooperatively binding cognate DNA sequence motifs residing in cis-regulatory elements (CREs), such as promoters and enhancers, and forming complexes that have regulatory potential to activate nearby genes (Kundaje et al., 2015; Reiter et al., 2017; Rubin et al., 2017). The quantitative changes in chromatin state and expression are hence highly dependent on the cis-regulatory code of motif patterns encoded in CREs (Arnosti et al., 1996; Banerji et al., 1981; Levo and Segal, 2014; Thanos and Maniatis, 1995). Previous studies have shown that the process of terminal differentiation alters the expression of thousands of genes, regulatory elements, proteins and metabolites (Michaletti et al., 2019). However, such efforts require increased temporal resolution to map dynamic regulation of subtle cell state transitions. While a number of regulators of epidermal differentiation have been previously identified (Hopkin et al., 2012; Lopez et al., 2009; Lopez-Pajares et al., 2015; Rubin et al., 2017; Segre et al., 1999; Sen et al., 2012; Truong et al., 2006), the combinatorial, dynamic cis-regulatory code of epidermal differentiation has remained elusive.

Recently, deep learning models such as convolutional neural networks (CNNs) have emerged as state-of-the-art predictive models of regulatory DNA. CNNs learn non-linear predictive functions that can accurately map DNA sequence to genome-wide profiles of regulatory activity by learning *de novo* predictive motif patterns and their higher-order combinatorial logic (Eraslan et al., 2019; Kelley et al., 2016, 2018; Zhou and Troyanskaya, 2015). Hence, interpreting these models could provide new insights into the cis-regulatory sequence code. We and others have recently developed powerful interpretation methods to extract rules of cis-regulatory logic from these black-box models (Avsec et al., 2020; Ching et al., 2018; Greenside et al., 2018; Shrikumar et al., 2017). These interpretable deep learning models have the potential to offer new insights into the cis-regulatory code of epidermal differentiation.

Here, we use a battery of assays to comprehensively profile the multi-modal landscape of chromatin and expression dynamics across a densely sampled time course of epidermal differentiation. We train robust CNN models that can accurately predict quantitative changes in chromatin accessibility from DNA sequence alone across the entire time course (**Fig. 1A**). We interpret the models to annotate tens of thousands of dynamic CREs with homotypic and heterotypic combinations of active motif instances then introduce an *in silico* combinatorial perturbation framework to decipher quantitative rules of cis-regulatory logic. We identify multiplicative and super-multiplicative effects of co-occurring motif combinations on chromatin accessibility and predict putative TFs that cooperatively bind these combinatorial motif patterns and link dynamic CREs to their putative target genes. Finally, we validate temporal dynamics and cis-regulatory logic of combinatorial motif rules on intrinsic regulatory activity across differentiation using massively parallel reporter assays (MPRAs). Genetic variants associated with diverse skin-related complex traits are found to be enriched in time-dependent combinatorial motif rules, supporting a potential disease-relevant role in mediating phenotypic effects. This integrative framework can be broadly applied to discover dynamic cis-regulatory logic across diverse cell states, cell types and conditions.

**Figure 1.**
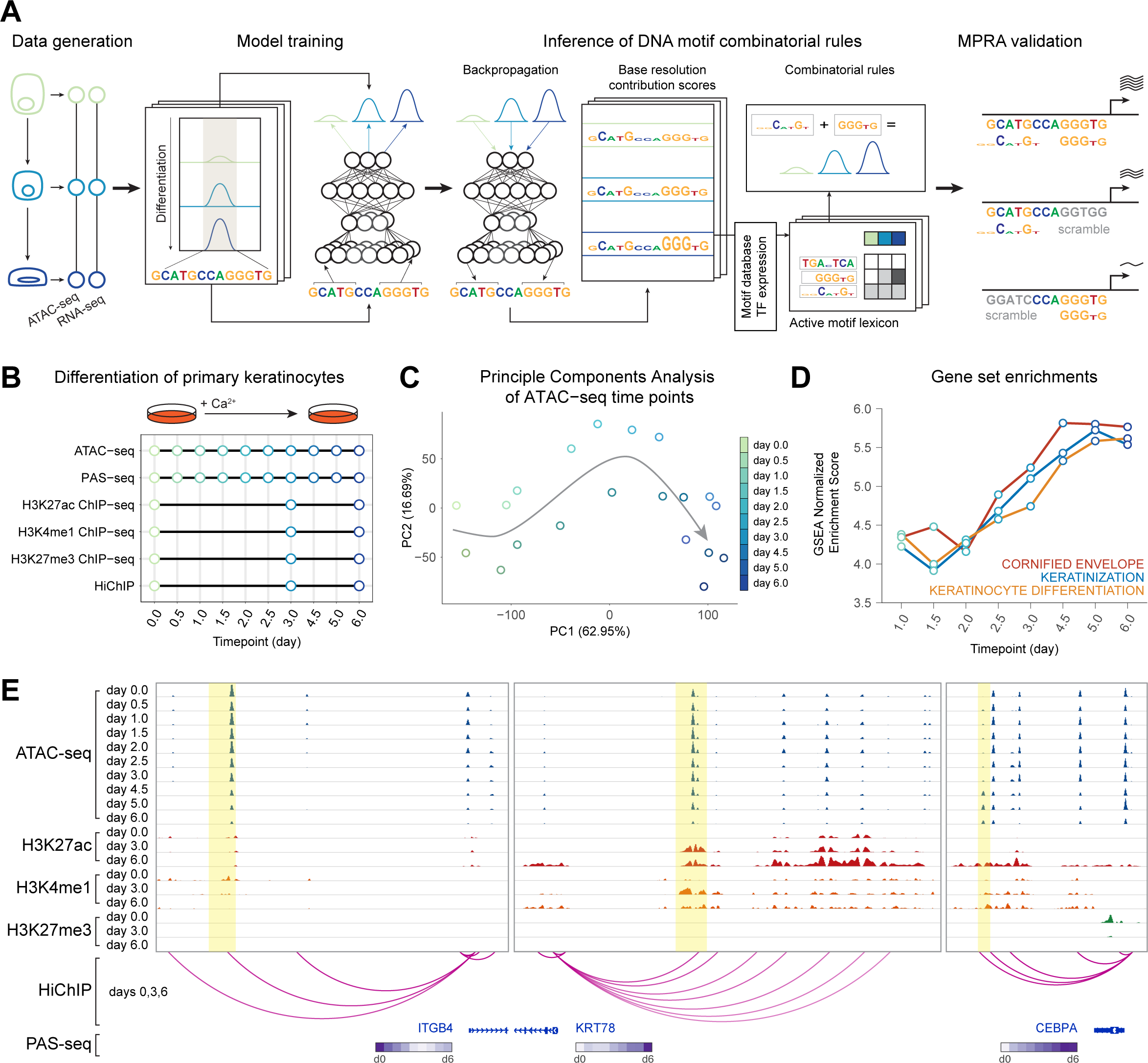
A high-resolution integrated multi-omic data resource in primary keratinocyte differentiation. (**A**) Schematic of the integrative framework for discovery of a dynamic, combinatorial cis-regulatory lexicon. Convolutional neural networks are trained to predict quantitative ATAC-seq signal from DNA sequence across a time course, augmented with prediction tasks for active chromatin marks. After model training, base-resolution contribution scores are inferred for all sequences using backpropagation-based interpretation methods, followed by motif scanning to identify predictive motif instances. *In silico* combinatorial perturbation analyses are used to identify interaction effects between co-enriched combinatorial motif rules. Gene expression (PAS-seq) across the time course enables identification of TFs that may bind motif rules and downstream target gene modules. Massively parallel reporter assays (MPRAs) validate predicted effects of combinatorial cis-regulatory logic. (**B**) Schematic of multi-omic data collected across the epidermal differentiation time course. (**C**) Principal component analyses of ATAC-seq data highlight time as the primary axis of variation. (**D**) Gene set enrichments validate veridical activation of keratinocyte differentiation in the gene expression data. (**E**) Representative loci around the *ITGB4, KRT78*, and *CEBPA* genes exhibit different trajectories of chromatin and expression dynamics.

## RESULTS

### An integrative resource profiling chromatin and expression dynamics during epidermal differentiation

To characterize the multi-modal regulatory landscapes of keratinocyte differentiation, transcriptional and chromatin state was profiled across multiple timepoints of calcium-induced *in vitro* differentiation (**Fig. 1B**) with high quality, replicated PAS-seq, ATAC-seq, H3K27ac ChIP-seq, H3K4me1 ChIP-seq, H3K27me3 ChIP-seq, CTCF ChIP-seq, and H3K27ac HiChIP experiments (**Fig. S1B-E, Tables S1-S5**). Principal component analysis (PCA) showed high consistency between biological replicates (**Fig. 1C, S1E**), and gene set enrichments validated veridical activation of keratinocyte differentiation in these data (Subramanian et al., 2005) (**Fig. 1D**). Important gene loci showed complex dynamic regulatory landscapes (**Fig. 1E**).

We used the ATAC-seq profiles to identify 225,996 high-confidence, reproducible CREs across all time points, of which 40,103 CREs exhibited significant variation of chromatin accessibility across the time course. Clustering these CREs based on their ATAC-seq profiles resulted in 15 distinct trajectories across differentiation (Love et al., 2014; McDowell et al., 2018) (**Fig. 2A**). Chromatin accessibility dynamics were strongly correlated with the dynamics of activating histone marks H3K27ac and H3K4me1. We associated the dynamic CREs to their putative target genes based on proximity and H3K27ac HiChIP data. Functional enrichment analysis of the gene sets associated with each dynamic CRE cluster revealed relevant and expected biological functions that were consistent with expression dynamics (**Fig. 2A, S1F,G**). For example, CREs linked to hemidesmosome genes, whose expression characterizes progenitors (Simpson et al., 2011), decreased in accessibility during differentiation. In contrast, CREs linked to differentiation genes (Candi et al., 2005) were enriched in trajectories activated during differentiation. Analysis of gene expression quantification from the PAS-seq experiments identified 3,069 dynamic RNA transcripts that clustered into 11 dynamic trajectories (**Fig. 2B**). Down-regulated genes encompassed progenitor proliferation genes and up-regulated genes including early and late epidermal differentiation genes. The dynamic CRE clusters and their associated target genes also exhibited synchronous concordance of gene expression and chromatin accessibility dynamics (**Fig. 2C,D**), consistent with a picture of coordinated waves of target gene activation driven by dynamically accessible CREs. These data map the dynamic regulatory landscape of keratinocyte differentiation and indicate a coordinated interplay of tens of thousands of regulatory regions with thousands of genes.

**Figure 2.**
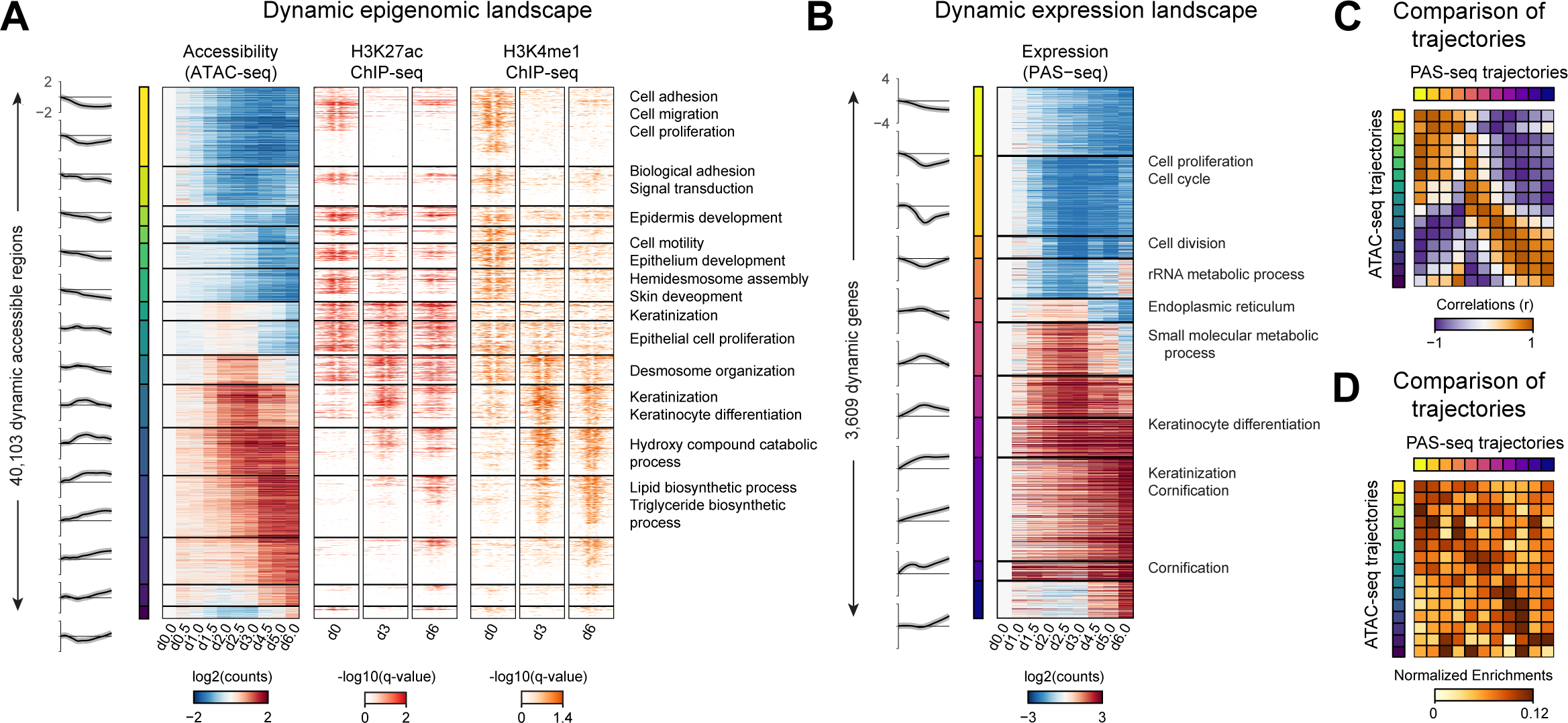
The dynamic chromatin and expression landscapes of epidermal differentiation. (**A**) ATAC-seq and ChIP-seq (H3K27ac, H3K4me1) heatmaps of 40,103 dynamic cis-regulatory elements, ordered by trajectories of dynamic accessibility; gene set enrichments of proximal genes (right). ATAC-seq signals are relative to day 0. (**B**) 3,609 dynamic genes identified in the same set of cells as in (A); gene set enrichments for each gene subset (right). RNA signals are relative to day 0. (**C**) Accessibility trajectories mapped to gene set trajectories. Correlation of accessibility trajectories (rows) to gene set trajectories (columns) by mean trajectory vectors. (**D**) Normalized enrichment of accessibility trajectory regions (rows) linked to each gene set (columns) when linked by proximity. Enrichments when linking with proximity show consistency with expected mappings by correlation.

### Predictive deep learning models of chromatin dynamics reveal a high-resolution cis-regulatory lexicon of epidermal differentiation

To learn predictive sequence models of chromatin dynamics, we trained multi-task CNNs to map 1 kb DNA sequences tiled across the genome to associated quantitative measures of ATAC-seq signal at 10 time points across the differentiation time course (**Fig. 3A**). We used a 10-fold chromosome hold-out cross-validation scheme to train and evaluate the predictive performance of the model (**Table S6**). We used a multi-stage transfer learning protocol. We first trained a reference model on large compendia of DNase-seq data from 431 diverse cell types and tissues. The reference model was then fine-tuned on the ATAC-seq data from our time course, accounting for the cross-validation structure of the 10 folds (**Fig. S2A, Tables S7, S8**). The model’s predictions on all held-out test set chromosomes were strongly correlated with the observed ATAC-seq signal in each time point (**Fig. 3A,B, Fig. S2B**). The model’s predictions for dynamic CREs were also strongly correlated with their measured ATAC-seq signal across the time course (**Fig. S2C**). This transfer learning approach substantially improved performance and stability of the model’s predictions across models and folds. The predictions of the models from the 10 folds for each CRE in each time point were subsequently calibrated and ensembled for downstream inference and prediction.

**Figure 3.**
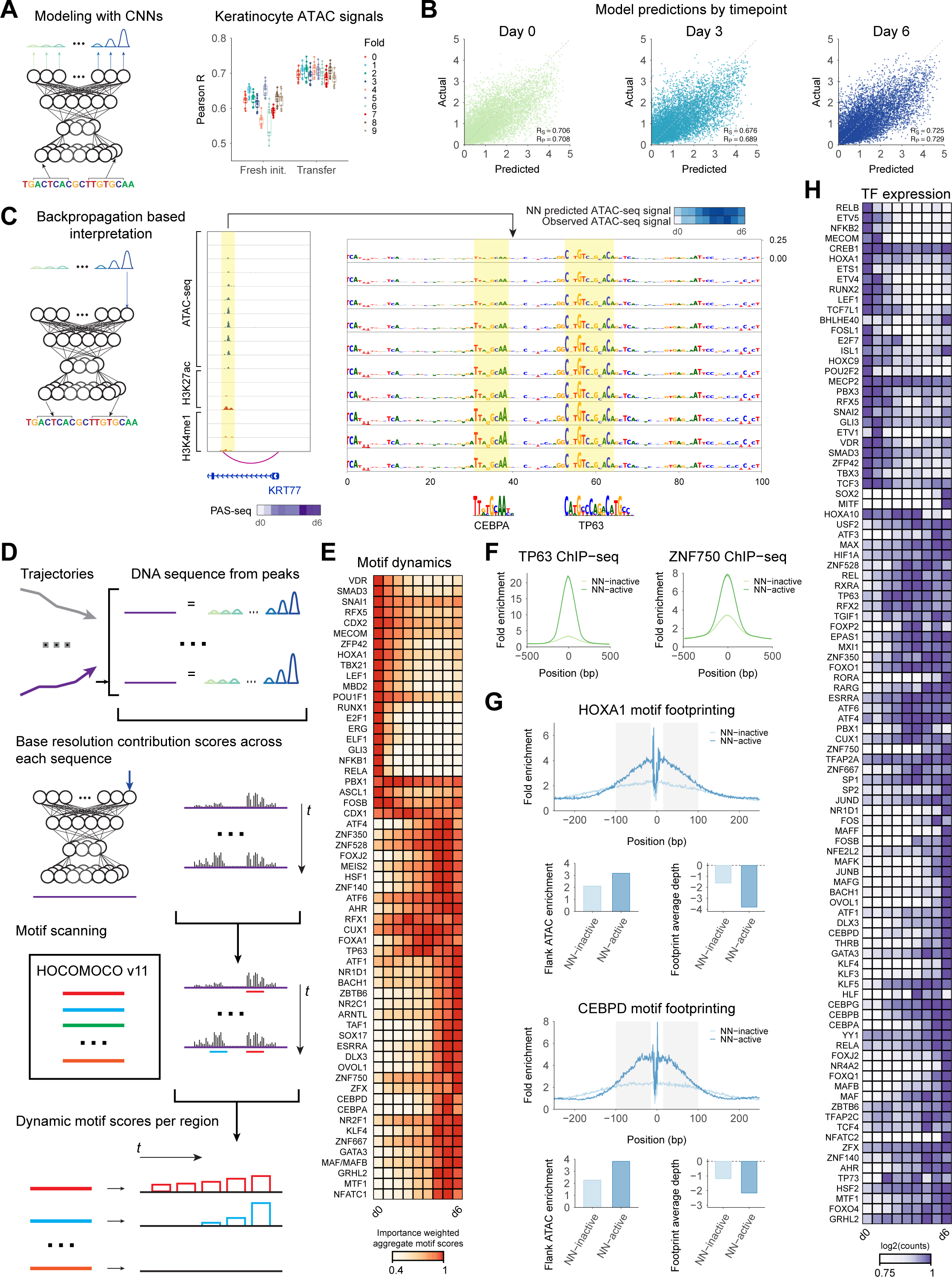
Deep learning models of chromatin accessibility reveal dynamic predictive motif instances across the differentiation time course. (**A**) Left: Schematic of a multi-task convolutional neural network that maps 1 Kb DNA sequences across the genome to quantitative chromatin accessibility signal across time points. Right: Pearson correlation (R) between predicted and observed accessibility across CREs of each time point for 10 folds of held-out test set chromosomes. (**B**) Scatter plots of predicted vs. observed accessibility signal (units of log depth normalized coverage) across CREs in test set chromosomes for three time points: (left to right) ATAC-seq at days 0, 3, and 6. (**C**) Left: Schematic of inference of base resolution contribution scores for a sequence with respect to predicted output at specific time points using efficient backpropagation methods. Right: A CRE linked via H3K27ac HiChIP to the promoter of the *KRT77* gene shows progressively increasing contributions of nucleotides in CEBPA and TP63 motifs across the time course in concordance with increasing accessibility. (**D**) Schematic for identification of predictive motif instances by scanning contribution scores with known motifs. Base-resolution contribution scores are inferred for all CREs corresponding to each dynamic trajectory. Contribution score weighted sequences are scanned and scored using a compendium of TF motifs (HOCOMOCO) to identify active, predictive motif instances. (**E**) Dynamics of predictive scores of motifs across time aggregated over all dynamic CREs. (**F**) Predictive motif instances of p63 and ZNF750 motifs exhibit higher ChIP-seq signal than predicted inactive motif instances in CREs. (**G**) ATAC-seq footprints are stronger at predictive motif instances of HOXA1 and CEBPD motifs relative to footprints at predicted inactive motif instances. (**H**) Dynamic expression patterns of TFs with correlated dynamics of matched predictive motifs across the differentiation time course.

Next, we used the ensemble of trained models to infer sequence features in each CRE that are predictive of chromatin accessibility at each time point. Specifically, we used efficient backpropagation methods (Shrikumar et al., 2017; Simonyan et al., 2014) that can infer contribution scores of each individual nucleotide in each input sequence with respect to the predicted output from the model at each time point (**Fig. 3C**). Although the sequence of a CRE is the same across all time points, the base-resolution contribution scores are dynamic and reflect the time-point specific activating or repressive effect of predictive sequence features through the lens of the model (**Fig. 3C**). To evaluate the potential functional consequences of predictive nucleotides highlighted by the model, we estimated the allelic imbalance of ATAC-seq reads (Harvey et al., 2015) of 16,686 single nucleotide polymorphisms (SNPs) in CREs. SNPs overlapping bases with high contribution scores were associated with larger allelic effect sizes (**Fig. S2D**). Additionally, model-derived predicted allelic effects using an *in-silico* mutagenesis approach were stronger for SNPs exhibiting statistically significant (FDR < 0.10) allelic imbalance than for SNPs that were allele-insensitive. These results indicate that the base-resolution contribution scores are enriched for nucleotides with putative functional effects on chromatin accessibility.

The base-resolution contribution scores highlighted short contiguous stretches of bases, reminiscent of TF binding motifs (**Fig. 3C**). Hence, to annotate predictive motif instances in each CRE in each time point, we scanned the base-resolution contribution scores with a non-redundant set of known sequence motifs of a large compendium of TFs (Kulakovskiy et al., 2018). We found 59 motifs with statistically significant (empirical *p* < 0.05) motif instances that were enriched in dynamic CREs (**Fig. 3D,E**). Dynamic, predictive motif instances inferred using this approach were strongly supported by matched TF occupancy and ATAC-seq footprints, indicating that they are likely capturing bound motif instances. For example, p63, ZNF750, and KLF4 ChIP-seq (Boxer et al., 2014; Liu et al., 2011; McDade et al., 2014) profiles exhibited higher occupancy at their predictive motif instances as compared to non-predictive motif instances within CREs (**Fig. 3F, S2E**). Similarly, ATAC-seq footprinting analysis (Li et al., 2019) identified stronger TF footprints at predictive motif instances (**Fig. 3G, S2F**). TFs belonging to the same family often bind very similar motifs. We mapped motifs to putative TFs that bind them by correlating predictive motif contribution scores across time points with mRNA expression of candidate TFs (**Fig. 3H, S2G-I, Table S9**). This analysis matched the 59 motifs to 100 putative TFs, many of which are known to be essential in keratinocyte differentiation, such as p63, CEBPA, GRHL2, AHR, FOSB, DLX3, VDR, ZNF750, MAFB, RARG, JUNB, KLF4, and OVOL1 (Boxer et al., 2014; Nair et al., 2006;Sen et al., 2012; Truong et al., 2006).

### Model interpretation enables discovery of combinatorial cis-regulatory logic of homotypic and heterotypic motif combinations

Next, we used the models to predict the quantitative effect of homotypic motif density and spacing on chromatin accessibility using synthetic DNA sequence inputs embedded with systematically varying density and spacing of motif instances of each of the 59 predictive motifs. While most TFs showed monotonic increases in accessibility with increasing homotypic motif density, some TFs displayed saturation effects indicating non-linear cooperative homotypic interactions (**Fig. S2J**). Increased spacing between homotypic motif instances was associated with decreasing accessibility up to 150 bp away (**Fig. S2K**) (Weingarten-Gabbay et al., 2019), indicating that combinatorial influence of homotypic TF binding on chromatin accessibility is constrained by proximity.

We then used two complementary *in-silico* motif perturbation analysis methods to quantify the influence of heterotypic pairs of co-occurring motifs on chromatin accessibility dynamics. The first approach quantifies the impact of *in silico* disruption of one instance of a predictive motif on the contribution scores of a co-occurring predictive instance of a different motif (Greenside et al., 2018). The second approach compares the sum of the marginal effects of *in silico* disruption of each motif instance to the effect size of jointly disrupting both motif instances on predicted chromatin accessibility (**Fig. 4A**). The models predict chromatin accessibility signal as the depth of normalized read coverage on log scale. Hence, additive effects on the log scale represent multiplicative effects on normalized read coverage. Motif pairs with joint effects larger than the sum of their marginal effects represent super-multiplicative interactions. Motif pairs whose joint effects are smaller than the sum of their marginal effects represent sub-multiplicative motif combinations that potentially act through independent, additive effects (**Fig. 4B, S3A**). We restricted heterotypic motif interaction analysis to motif pairs with enriched co-occurrence of predictive motif instances in the dynamic CREs (**Fig. 4C**). Co-occurrence statistics using only predictive motif instances instead of all motif instances revealed more specific and less promiscuous motif pairs (**Fig. S3B,C**). For each heterotypic pair of motifs, we estimated *in silico* interaction effects for all dynamic CRE sequences containing predictive instances of both motifs. A majority of the enriched co-occurring motif pairs exhibited multiplicative (log-additive) and super multiplicative effects (**Fig. 4D**), indicating extensive cooperativity between co-binding TFs through heterotypic motif syntax.

**Figure 4.**
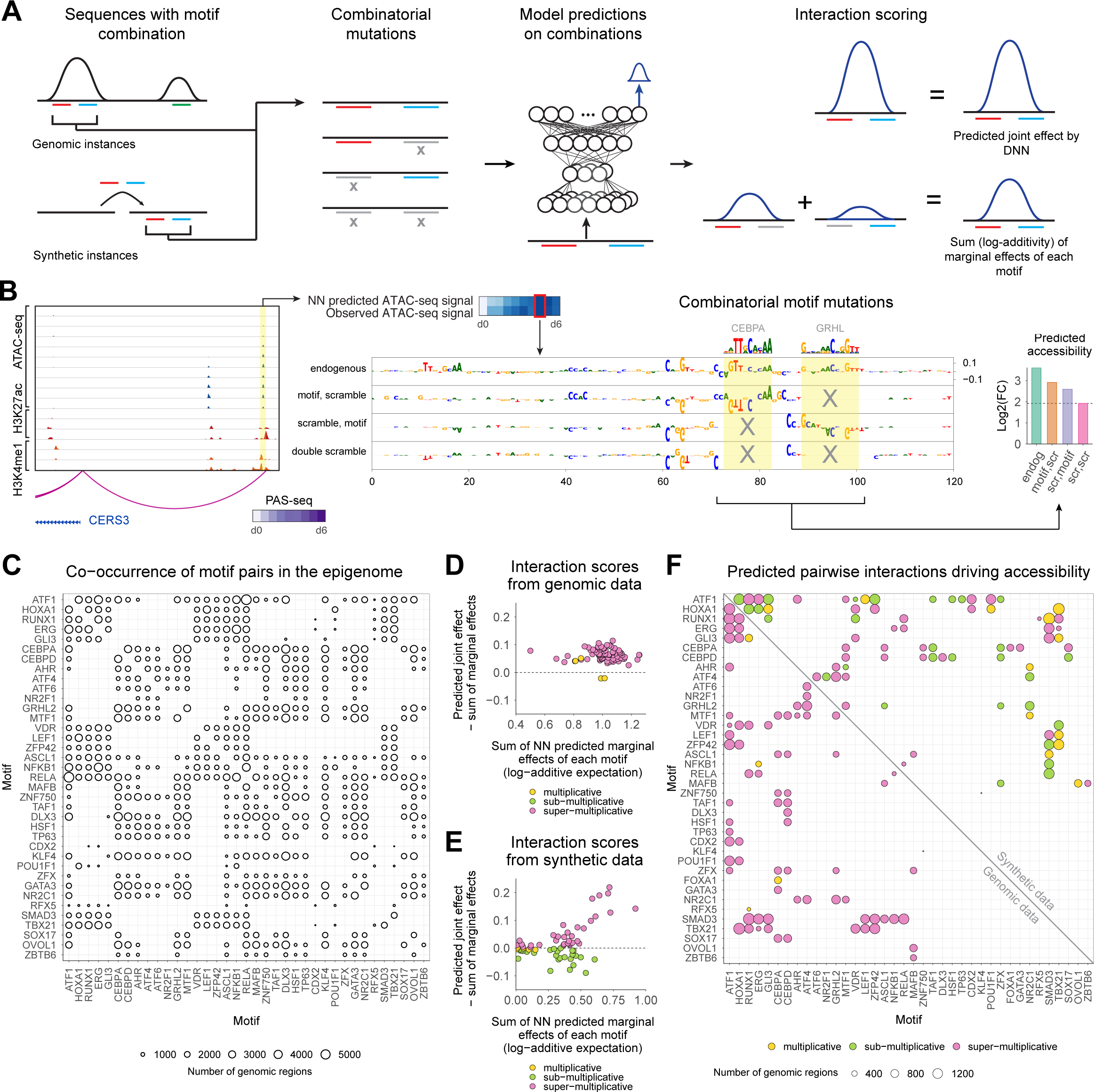
Combinatorial *in silico* perturbation analysis to infer heterotypic cis-regulatory logic. (**A**) Schematic for combinatorial *in silico* perturbation analysis. All genomic instances of CREs containing significantly co-occurring motif pairs are evaluated. Motif pairs are also embedded in synthetic scrambled background sequences for orthogonal evaluation. For each candidate sequence containing a motif pair, the neural network is used to predict changes in chromatin accessibility due to marginal perturbation of each motif and joint perturbation of both motifs. The joint effects are compared to the sum of the marginal effects (log-additivity) to test super-multiplicative, multiplicative (log-additive) or sub-multiplicative joint effects. (**B**) Example locus around the *CERS3* gene. A CRE that loops to the *CERS3* promoter contains active motif instances of CEBPA and GRHL motifs. Perturbing either of the motifs impacts contribution scores of the partner indicating an interaction effect. The contribution score tracks from top to bottom are the endogenous sequence, the sequence with GRHL motif scrambled, the sequence with CEBPA scrambled, and the sequence with both scrambled (double scramble). The right plot shows the predicted accessibility for the endogenous sequence, sequences with marginal perturbations of individual motifs and joint perturbations (as the baseline). The motifs exhibit a multiplicative (log-additive) joint effect. (**C**) Number of CREs supporting significantly co-occurring predictive pairs of motifs. (**D**) Scatter plot comparing the difference between the joint effect on predicted accessibility and the sum of the predicted marginal effects (y-axis: neural net predicted joint effect minus the sum of the marginal effects) to the sum of marginal effects (x-axis) of motif perturbations for all significantly co-occurring motif pairs using genomic sequences. Super-multiplicative pairs (pink) fall above the dashed line, multiplicative pairs (yellow) fall near and on the dashed line and sub-multiplicative (green) pairs fall below the dashed line. (**E**) Scatter plot comparing the difference between the joint effect on predicted accessibility and the sum of the predicted marginal effects (y-axis) to the sum of marginal effects (x-axis) of motif perturbations for all significantly co-occurring motif pairs using synthetic sequences. (**F**) Comparison of interaction effects of all significantly co-occurring motif pairs that exhibit skin-related functional enrichments using genomic sequences (below diagonal) and synthetic sequences (above diagonal). The size of the circles represents the number of dynamic CREs supporting each motif pair. The color of the circle represents the type of interaction: super-multiplicative (pink), multiplicative (yellow), sub-multiplicative (green).

We also computed *in silico* interaction effects for motif pairs after embedding them in synthetic scrambled background sequences to avoid cryptic cooperative effects induced by additional predictive motifs in the endogenous context (**Fig. 4E**). We observed more sub-multiplicative motif interactions in these synthetic backgrounds as compared to endogenous sequence context. These differences indicate that the native genomic context likely encodes higher-order cooperative interactions between the tested motif pairs and additional motif partners. In order to winnow down the motif pairs to those with likely functional roles, we computed enrichments of functional terms using proximal gene sets associated with all CREs harboring predictive instances of each motif pair (**Fig. S4A**) and restricted to those that were enriched for skin related functional terms (**Fig. 4F**). We thus obtained a core lexicon comprising 80 heterotypic pairs of significantly co-occurring TF motifs linked to distinct processes at different stages of epidermal differentiation.

This combinatorial lexicon implicates known and novel cooperative partners (**Fig. S4A**). ZNF750 motif was found to strongly interact with CEBP family member motifs (CEBPA/CEBPD), both of which are known to be important in KRT10 regulation (Maytin et al., 1999). The ATF1 motif is present in stem cell maintenance rules, such as ATF1 with GLI1, as well as late differentiation rules, such as ATF1 with TP63. Notably, of the TFs that can bind to the ATF1 motif, CREB1 is most expressed at the beginning and end of differentiation while ATF1 increases in expression. NFKB/REL motifs are only present in stem cell maintenance rules, supporting a role for NFKB/REL motifs in progenitor state maintenance (Li et al., 2000). Notably, of the TFs that bind to the NFKB/REL motifs, RELB/NFKB2 decrease in expression while REL/RELA increase in expression. These rules in conjunction with matched TF dynamics demonstrate that precise targeting of gene modules and coordination of activation and deactivation relies on combinatorial sequence and specific TF family member expression.

### Massively parallel reporter experiments validate temporal dynamics and cis-regulatory logic of the combinatorial motif lexicon

Next, we validated the temporal dynamics and the quantitative effects of the combinatorial motif lexicon on intrinsic regulatory potential using massively parallel reporter (MPRA) experiments in multiple time points of *in vitro* differentiation. We used the model-derived, predictive motif annotations of all dynamic CREs to design libraries for the MPRA experiments. We designed 160 bp constructs for 19 randomly selected endogenous CRE sequence examples from each of the 80 heterotypic motif pairs, mutants with combinatorially scrambled motif instances (individually and jointly) as well as corresponding positive and negative controls for the MPRA library – a total of 77,090 sequences (**Fig. 5A, S5A,S5B, Table S10**). MPRA read outs for the entire library were obtained on days 0 (progenitor state), 3 (early differentiation), and 6 (late differentiation) of the differentiation time course (**Fig. S5C,D,E**). Sample clustering and principal component analysis demonstrated high reproducibility and clear separation between the progenitor state at day 0 and the differentiated state at days 3 and 6 (**Fig. S5F**).

**Figure 5.**
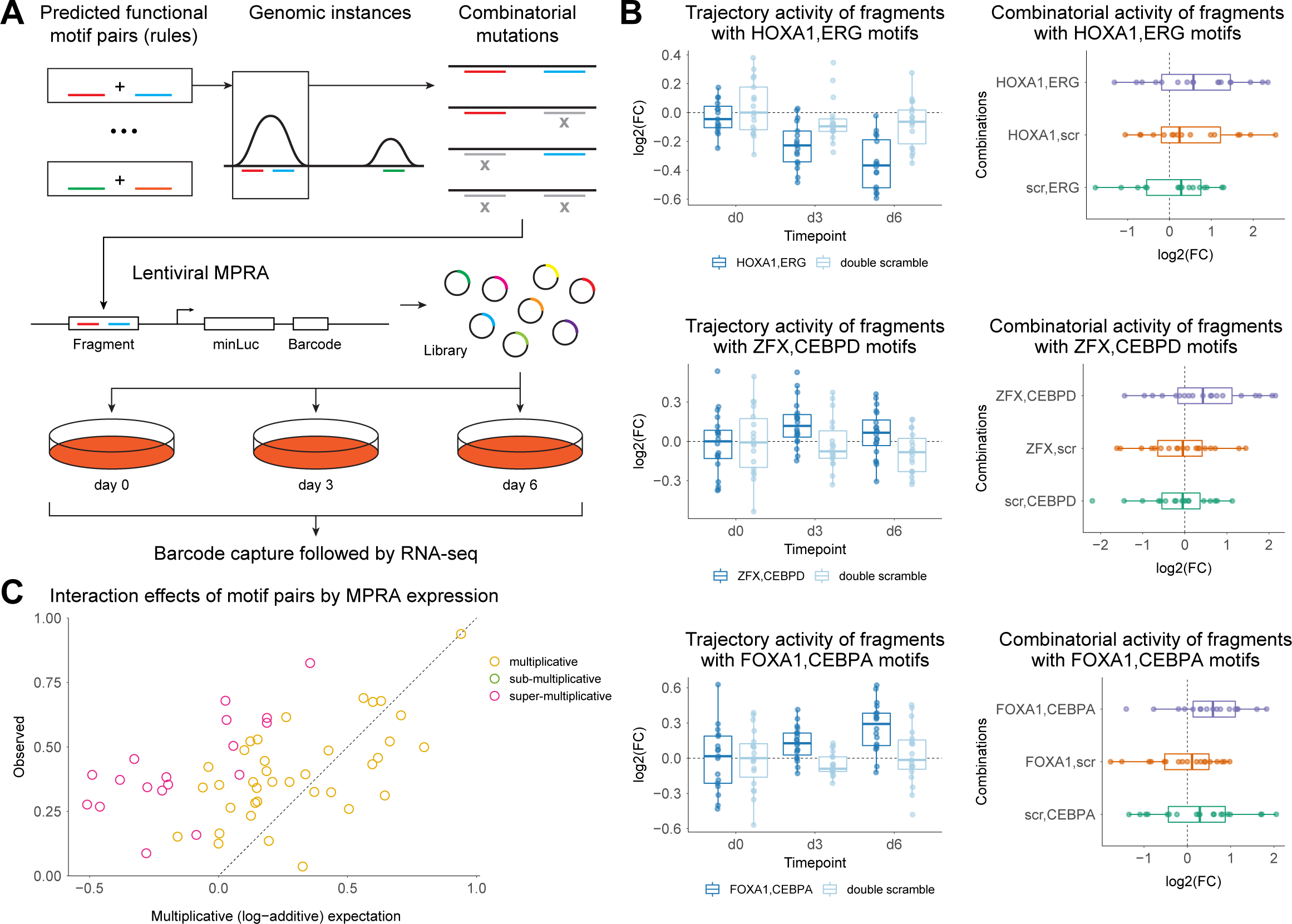
Validation of combinatorial motif pairs using massively parallel reporter assays. (**A**) MPRA design. For each of the derived combinatorial rules, genomic instances of each rule were selected randomly and the motif pair in the instance was combinatorially scrambled. All combinatorial versions of the sequence were added to the MPRA library, which was lentivirally inserted into primary keratinocytes. These cells were induced to differentiate, and reporter RNA was collected at days 0, 3, and 6, (**B**) Examples of three combinatorial rules: HOXA1/ERG motif pair (progenitors), ZFX/CEBPD motif pair (early differentiation), and FOXA1/CEBPA motif pair (late differentiation). Left column plots show observed expression across time for the endogenous sequences as well as double motif scrambled mutants, normalized to day 0. Right column shows the combinatorial dynamics of genomic instances of each rule, relative to joint motif scrambled mutants. (**C**) Summary of combinatorial interaction effects of all temporally valid motif pairs. The scatter plot compares the joint effect (log fold change of reporter expression) of each motif pair (y-axis) relative to the sum (log-additivity) of the marginal effects of each motif. Motif pairs with significant super-multiplicative effects on fold-change are marked in pink above the diagonal. Sub-multiplicative (log-additive) motif pairs are marked in yellow. No motif pairs were found to have sub-multiplicative effects.

First, we compared the MPRA-measured expression for the endogenous genomic regulatory sequences in our library to their corresponding measured and predicted ATAC-seq signal as well as H3K27ac signal in matched time points. We observed low correlation between MPRA expression and observed ATAC-seq signal (Pearson Ρ = 0.097, Spearman Ρ_#_ = 0.075), predicted ATAC-seq signal (Pearson Ρ = 0.088, Spearman Ρ_#_ = 0.065) and observed H3K27ac signal (Pearson Ρ = 0.061, Spearman Ρ_#_ = 0.046) (**Fig. S5G**), suggesting fundamental differences between the MPRA-derived intrinsic measures of regulatory potential and endogenous chromatin state of regulatory sequences. However, we found that simple linear models that used the non-linear sequence representation encoded in the final layer of the ATAC-seq CNN models as inputs were able to fit the MPRA expression levels with improved correlation (Pearson Ρ = 0.344, Spearman Ρ_#_ = 0.3) (**Fig. S5G**). These results suggest that the combinatorial sequence features that are predictive of ATAC-seq signal are also predictive of MPRA activity after a simple linear transformation. Hence, we postulated that the MPRAs could be used to validate the different types of motif interactions discovered by the models trained on the ATAC-seq data.

Since we observed synchronicity between chromatin dynamics and expression dynamics of associated putative target genes, we considered a heterotypic pair temporally valid if it produced a synchronous effect in reporter expression compared to the measured and predicted chromatin accessibility dynamics of the CREs containing the pair. For example, for tested sequences containing the HOXA1-ERG motif pair, reporter activity decreases during differentiation, synchronous with the accessibility dynamics of the CREs containing this pair (**Fig. 5B, S5H**). Using this criterion, 55 of the 80 heterotypic motif pairs (68%) were validated for temporal dynamics. Of these, 43 of the pairs (78%) showed significant differential activity relative to the mutated constructs in which both motifs were scrambled, suggesting that these motif pairs are key drivers of regulatory potential for these CREs. Next, we used the combinatorially scrambled mutant sequences to determine whether the heterotypic motif pairs had multiplicative, super-multiplicative or sub-multiplicative effects on reporter expression. Of the 55 temporally valid motif pairs, we found that 18 pairs had super-multiplicative effects, 37 rules had multiplicative (log-additive effects) and none of the rules exhibited sub-multiplicative effects on reporter expression (**Fig. 5C, 6A**). Hence, the MPRA experiments support the multiplicative and super-multiplicative cooperative effects of motif pairs on chromatin accessibility as predicted by the model.

### The cis-regulatory lexicon of epidermal differentiation is enriched for skin disease-associated genetic variation

Utilizing imputed GWAS studies from the UK Biobank database (http://www.nealelab.is/uk-biobank/), we observed that 493 genome-wide significant variants for a curated set of skin phenotypes were found in 295 CREs (from 2,092 total genome-wide significant variants across the phenotypes). These phenotypes included a variety of human skin diseases characterized by dysregulated epidermal differentiation, such as pre-malignant actinic keratoses, dermatitis, psoriasis, rosacea, and acne vulgaris. To test whether the combinatorial motif lexicon was enriched for non-coding variants associated with these complex skin phenotypes, we used LD-score regression (Finucane et al., 2015, 2018) in conjunction with the curated UK Biobank phenotypes and curated GWAS studies with summary statistics (Hirata et al., 2018; Paternoster et al., 2015). Genetic variants associated with skin-related diseases and traits were enriched in CREs containing specific motif rules with distinct temporal activity (**Fig. S6A**), indicating that disruption of cooperative TF interactions that regulate epidermal differentiation could mediate disease risk and pathogenesis. No such enrichment was observed in variants linked to other traits, such as body mass index (BMI). Skin disease variants were enriched in lexicons linked to in a manner consistent with pathogenic features of the corresponding skin disease. For example, motif pairs that influence the late stages of differentiation were enriched for heritability associated in acne, which is pathogenically linked to abnormal terminal follicular keratinization. In contrast, motif pairs that influence the mid stages of differentiation were enriched in variants linked to heritability for actinic keratosis, a pre-malignant neoplasm characterized by a failure to fully enter the normal differentiation pathway. These results suggest heterogeneous effects of different components of the cis-regulatory motif lexicon on different phenotypic outcomes through different stages of differentiation relevant to pathogenesis of human disease.

## DISCUSSION

Here, we present a unique resource for deciphering the cis-regulatory code of epidermal differentiation. Dense longitudinal profiling of the transcriptome and epigenome through the differentiation process enabled the identification of distinct dynamic trajectories of 40,103 dynamic CREs driving synchronous changes in gene expression of linked target genes. The depth and breadth of the data allowed training of deep learning models to infer the combinatorial lexicon of cooperative transcription factor binding sites encoded in the dynamic CREs at single base resolution. Massively parallel reporter experiments validated predicted temporal dynamics and cis-regulatory logic involving cooperative TF interactions.

This integrative resource serves as a repository of hypotheses about combinatorial cis-regulatory control of several key processes in epidermal differentiation (**Fig. 6B**). We find a progenitor maintenance lexicon including RELB, NFKB2, ETS1, SMAD3, and RUNX1 motifs that jointly orchestrate deactivation and disassembly of hemidesmosomes, structural proteins that anchor keratinocytes to the basement membrane. Notably the associated TFs decrease in expression quickly, within the first 12 hours of initiating differentiation. The deactivation of this lexicon thus enables prompt migration of cells from the basement membrane. We also identified intricate interplay of motifs in an early differentiation lexicon involving ATF4, ATF6, GRHL2, MTF1, and NR2C1 motifs that associates with induction of early differentiation genes. ATF4 motif, the hub motif in this lexicon, is bound by ATF4 whose mRNA is induced earlier, at day 2.5, while the other TFs follow, suggesting that ATF4 may be responsible for priming the landscape for subsequent activation of specific gene modules directed by its partners. ATF4 is also the first TF to decrease in expression later in the time course around day 5, suggesting that ATF4 leaving the landscape might bookend this phase of differentiation. In late differentiation, we discovered a lexicon comprising HSF2, CEBPD, ZFX, CEBPA, and ZNF750 motifs that regulate a module of genes involved in fatty acid catabolism, an essential process for cornification and maintenance of skin barrier function. ZNF750 is one of the last TFs to sharply increase in expression around day 5.5, consistent with the essential role of ZNF750 in orchestrating terminal skin barrier formation (Birnbaum et al., 2006; Sen et al., 2012). Finally, the enrichment of skin disease associated variants in specific rules of the cis-regulatory lexicon suggest that this approach could prove useful in future efforts aimed at fine mapping causal variants and genes as well as providing mechanistic insights into how these variants might disrupt key pathways in skin differentiation.

**Figure 6.**
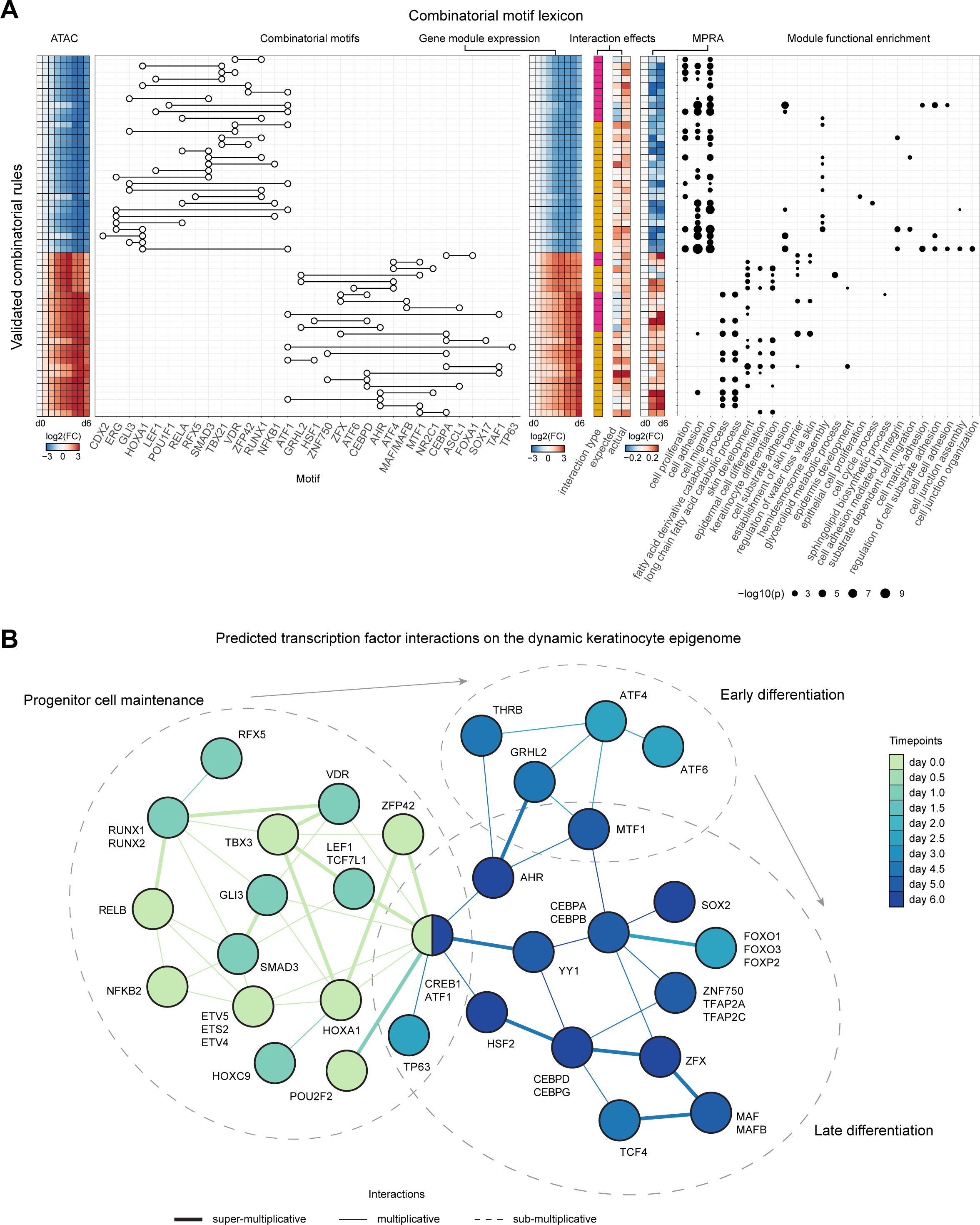
A combinatorial motif lexicon in keratinocyte differentiation. (**A**) Summary of the validated combinatorial lexicon of motif pairs. Left to right: heat map of ATAC-seq dynamics associated with each motif pair; motif pairs (each row is a distinct motif pair); expression dynamics of putative downstream target genes associated with each motif pair; type of interaction (pink: super-multiplicative, yellow: multiplicative), expected sum of marginal effects compared to joint effects in the MPRA; enriched functional terms for downstream target gene sets associated with each motif pair. (**B**) Predicted TF interactions on the dynamic keratinocyte epigenome. Each node is a TF (or multiple) that are matched to their DNA binding motifs, and the node is filled in by the most active timepoint by TF expression. Each edge is a predicted interaction on regulatory DNA, as predicted by the validated motif interactions from MPRA, between the TFs. Each edge is colored by the most active timepoint by accessibility. Edges are weighted by the predicted interaction effect: super-multiplicative, multiplicative, or sub-multiplicative.

The cis-regulatory code is more than the sum of its parts. The interpretable, deep learning framework presented here (https://github.com/kundajelab/tronn) provides a generalizable approach to move beyond static catalogs of cis-regulatory “parts-lists” (Bentsen et al., 2020; ENCODE Project Consortium, 2012; ENCODE Project Consortium et al., 2020; Fornes et al., 2020; Kulakovskiy et al., 2018; Kundaje et al., 2015; Luo et al., 2020; Vierstra et al., 2020; Weirauch et al., 2014) to predictive, quantitative models of higher-order cis-regulatory logic. Previous advances in deep learning model interpretation methods have largely focused on discovering motif representations, active motif instances and their co-occurrence patterns (Alipanahi et al., 2015; Ghandi et al., 2014; Kelley et al., 2016, 2018; Maslova et al., 2019; Sanford et al., 2020; Zhou and Troyanskaya, 2015). The current *in-silico* combinatorial perturbation framework extends this to enable discovery of quantitative rules of homotypic and heterotypic cis-regulatory logic such as the multiplicative and super-multiplicative effects of frequently co-occurring motif combinations on chromatin accessibility. Unlike previous studies that have investigated the critical regulatory role of cooperative TF binding in limited contexts, this approach allows comprehensive, genome-wide elucidation of these effects, at the resolution of individual CREs in dynamic processes such as cellular differentiation.

The present analyses also reconcile the influence of cis-regulatory logic on endogenous chromatin state and intrinsic regulatory potential. MPRAs offer a powerful experimental platform to test the effects of motif combinations on reporter gene expression activity (Sharon et al., 2012; Smith et al., 2013). However, interpretation of MPRAs designed to test endogenous properties of regulatory DNA is challenging since the sequences are tested outside their native genomic context. We show that the sequence features learned by deep learning models of chromatin accessibility, however, are also predictive of MPRA read-outs. Hence, despite the fundamental differences between these assays, we observe largely consistent rules of cooperative cis-regulatory logic derived from predictive models of chromatin state and the MPRA experiments.

Our study has several limitations, providing scope for future enhancements. Our cis-regulatory lexicon is biased towards activators due to inherent biases of our chosen assays and our modeling focus on active CREs in each time point. Models trained on markers of repression as well as differential effects across time points could reveal cis-regulatory sequences associated with dynamic repression. The current work also does not model the combinatorial effects of multiple CREs on gene expression. However, our chromatin-based models serve as a foundation for higher-order predictive models of gene expression which could be interpreted using similar *in silico* combinatorial perturbation strategies to decipher the distributed cis-regulatory code. We do not directly model the combinatorial influence of trans-acting factors on chromatin state and gene expression. Future modeling efforts, however, could be designed to jointly learn cis and trans regulatory logic from multimodal perturbation experiments (Sanford et al., 2020). Finally, extensions of these models to continuous cell state transitions from multi-modal single cell readouts of chromatin state and expression are exciting avenues for future research.

## ACKNOWLEDGMENTS

We thank members of the Kundaje, Greenleaf, and Khavari laboratories for discussions and Stanford Computing (Sherlock) and Pacific Computing Consortium (Nautilus) for computing resources. This work was supported by the USVA Office of Research and Development I01BX00140908 (P.A.K.), NIH CA142635, AR45192, AR076965 and HG007919 (P.A.K),1DP2GM123485 (A.K.), RM1-HG007735 (H.Y.C.). H.Y.C. is an Investigator of the Howard Hughes Medical Institute. D.S.K, A.K., and P.A.K. conceived the project.

## AUTHOR CONTRIBUTIONS

D.S.K, A.K., W.J.G., and P.A.K. conceived the project. D.S.K., V.R., D.R., J.C., A.J.R., N.J., L.D.,A.K., M.S., Z.Z. performed experiments and performed analysis. A.K. and P.A.K. guided experiments and data analysis. D.S.K, W.J.G., P.A.K. and A.K. wrote the manuscript with input from all authors.

## SUPPLEMENTAL FIGURE LEGENDS

**Figure S1.**
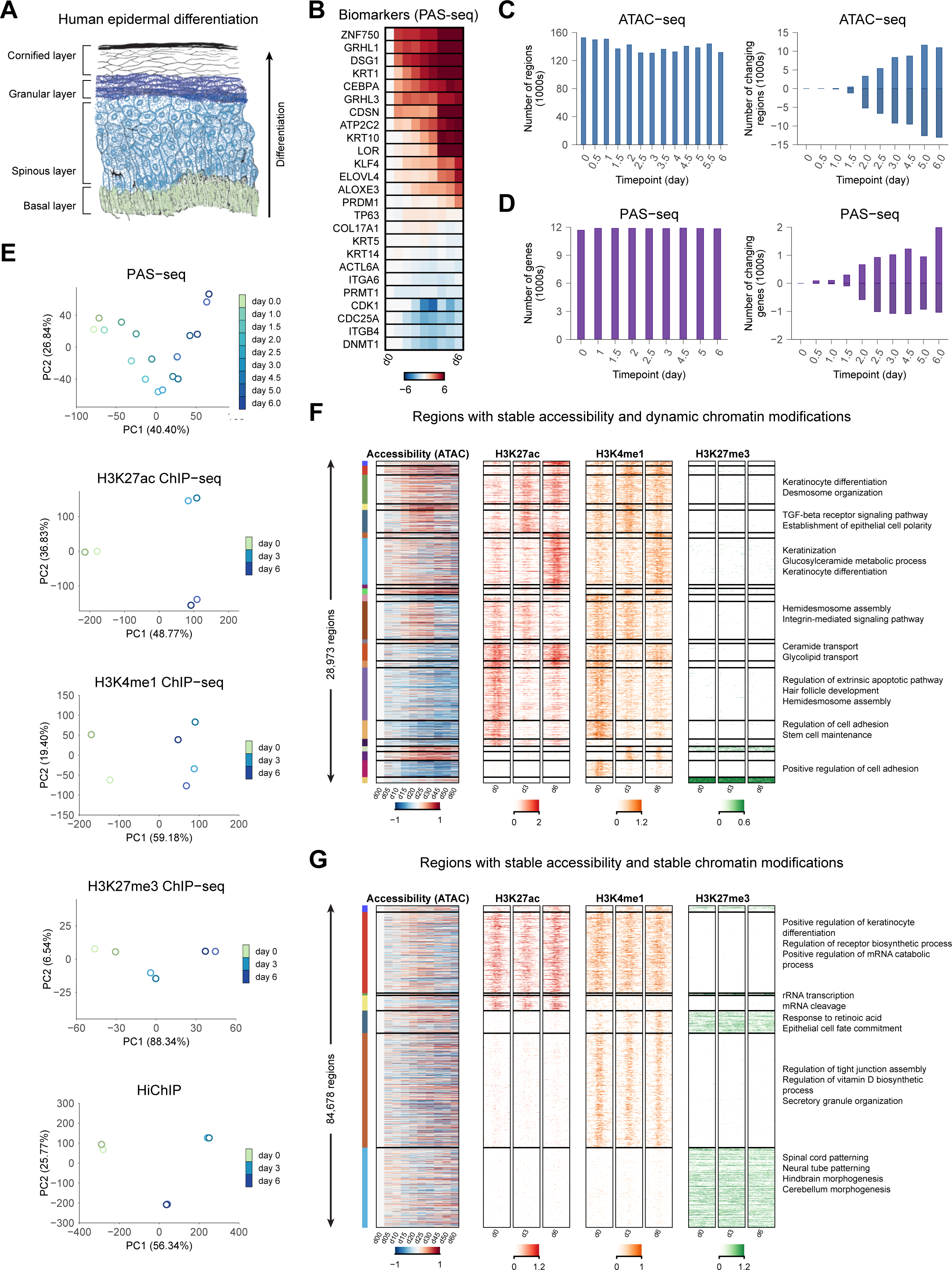
Data quality and other characteristics of the regulatory landscape. (**A**) Morphology schematic of normal human epidermis. (**B**) Selected biomarker gene panel from PAS-seq, demonstrating proper differentiation across time in vitro. (**C**) Global statistics on ATAC-seq. Left plot shows the number of reproducible peaks across the timepoints. Right plot shows the number of up and down regulated differential peaks across time, using day 0 as the background. (**D**) Global statistics on PAS-seq. Left plot shows the number of expressed genes (> approximately 1TPM) at each timepoint. Right plot shows the number of up and down regulated differential genes across time, using day 0 as the background. (**E**) PCA of other datasets: ATAC-seq, H3K27ac ChIP-seq, H3K4me1 ChIP-seq, H3K27me3 ChIP-seq, HiChIP. (**F**) Analysis of regions with stable (invariant) accessibility and dynamic chromatin modifications surrounding them (28,973 regions). The regions are clustered according to their dynamic chromatin mark patterns and marked with enriched GO terms accordingly. (**G**) Analysis of regions with stable (invariant) accessibility and stable chromatin modifications (84,678 regions). The regions are clustered according to combinatorial chromatin states and marked with enriched GO terms accordingly.

**Figure S2.**
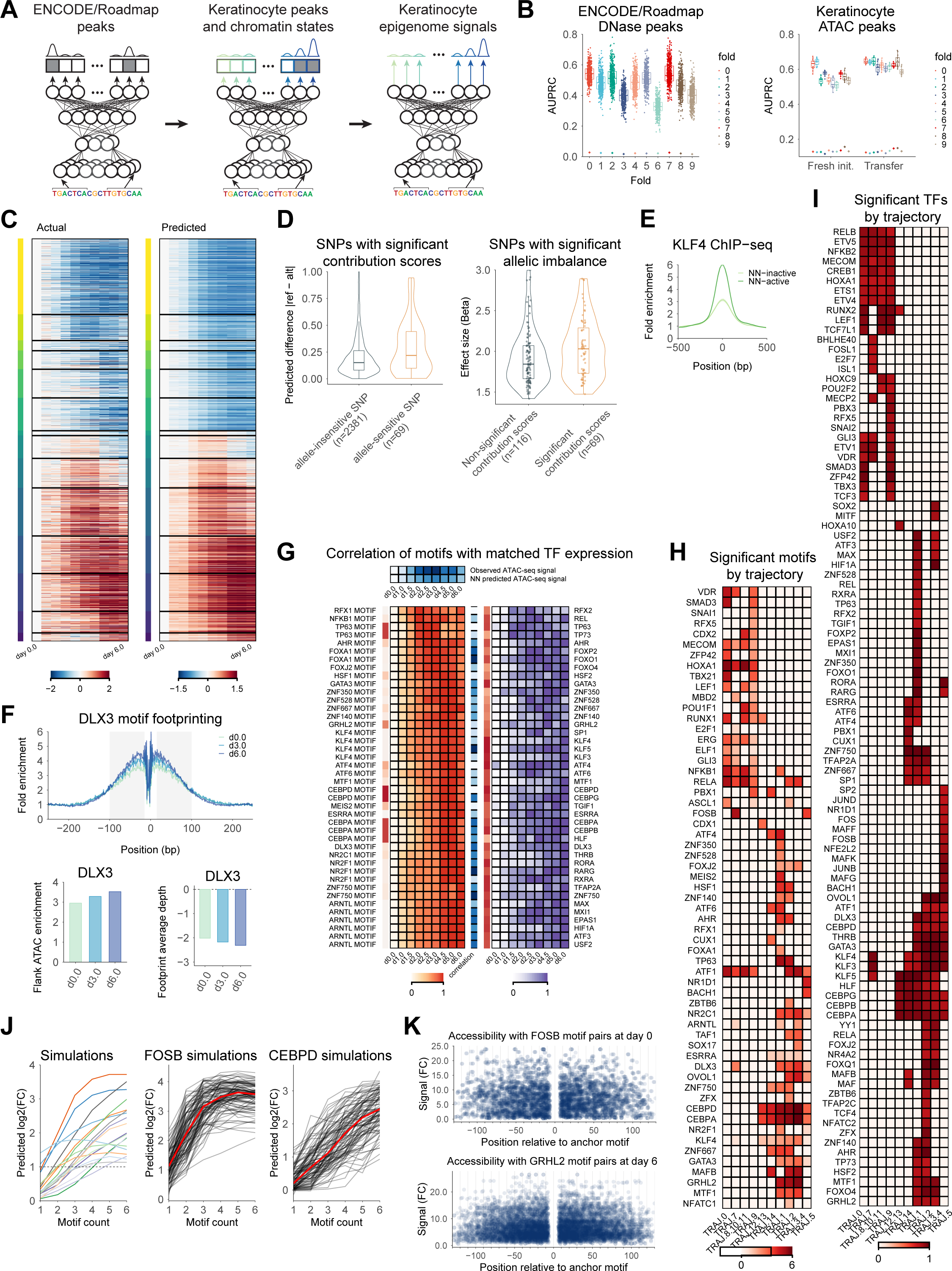
Extended analysis of deep neural net models and their utility for biological interpretation. (**A**) Schematic describing transfer learning set up. From left to right: first, models are trained on a large compendium of DNase-seq and TF ChIP-seq datasets from ENCODE and Roadmap, predicting accessibility and TF binding from DNA sequence; the weights from these models are then used as initialization weights for a classification model with keratinocyte-specific datasets from this study to predict keratinocyte accessibility/histone marks/trajectories from DNA sequence; finally, the weights from the classification model are used as initialization weights for a regression model in which DNA sequence is mapped to accessibility, H3K27ac, and H3K4me1 signals. This operation is done with 10-fold cross validation, where a test set is held out for the entire transfer learning process. (**B**) Metrics across transfer learning. Left plot shows the AUPRC for the ENCODE/Roadmap classification tasks (DNase and TF ChIP-seq) across 10 folds. Right plot shows AUPRC for accessibility in keratinocyte timepoints across 10 folds, considering transfer learning or fresh initialization (random seeded weights). (**C**) Heatmaps of observed ATAC signal vs neural net predicted ATAC signal across dynamically accessible regions. (**D**) Validation of contribution scores by comparing to SNPs exhibiting significant allelic imbalance of ATAC-seq signal. Left: Comparison of effect sizes of allelic imbalance of ATAC-seq signal, between SNPs overlapping non-significant contribution scores and those overlapping significant contribution scores. Right: comparison of model derived allelic effect predictions (reference allele - alternate allele) on SNPs overlapping significant contribution scores, separated by whether the SNP was considered allele-sensitive (FDR < 0.10) or not allele-sensitive. (**E**) Predictive, active motif instances of KLF4 show higher ChIP-seq signal relative to inactive motifs in CREs. (**F**) Footprinting across time demonstrates dynamic footprinting. DLX3 motif is shown here, where the accessibility increases across time while the footprint also deepens across time. (**G**) Example map of motif contribution scores across time points and expression of matched TFs for one trajectory pattern. Top: comparison of observed ATAC signal with the neural net predicted ATAC signal. Left heatmap: Motif contribution scores across timepoints. Right heatmap: Expression patterns of matched TFs (TFs that bind the motif and exhibit expression dynamics that is concordant with motif dynamics) across timepoints. Each row represents a matching expressed TF to the motif in the left heatmap row. (**H**) Heatmap showing which motifs were found significant in which trajectory patterns. (**I**) Heatmap showing which TFs were found with correlating patterns (r > 0.8) to their matching PWMs and which trajectory pattern. (**J**) Analysis of homotypic density on chromatin accessibility using synthetic sequences. Synthetic scrambled background sequences were embedded with varying number and spacing of motif instances of each predictive motif. The neural network was used to predict chromatin accessibility. Left: Each curve summarizes the predicted accessibility with increasing motif density for each motif averaged over multiple synthetic backgrounds. Middle/right: Predicted chromatin accessibility for increasing density of FOSB, and CEBPD motifs. Each black curve represents a specific random synthetic background sequence, while the red curve is the average pattern across all backgrounds. (**K**) Predicted chromatin accessibility as a function of distance between two instances of a motif embedded in synthetic backgrounds. Top: accessibility for FOSB motif pairs, showing decreasing accessibility when the FOSB motifs are farther apart. Bottom: accessibility for GRHL2 motif pairs, showing decreased accessibility when the GRHL2 motifs are farther apart.

**Figure S3.**
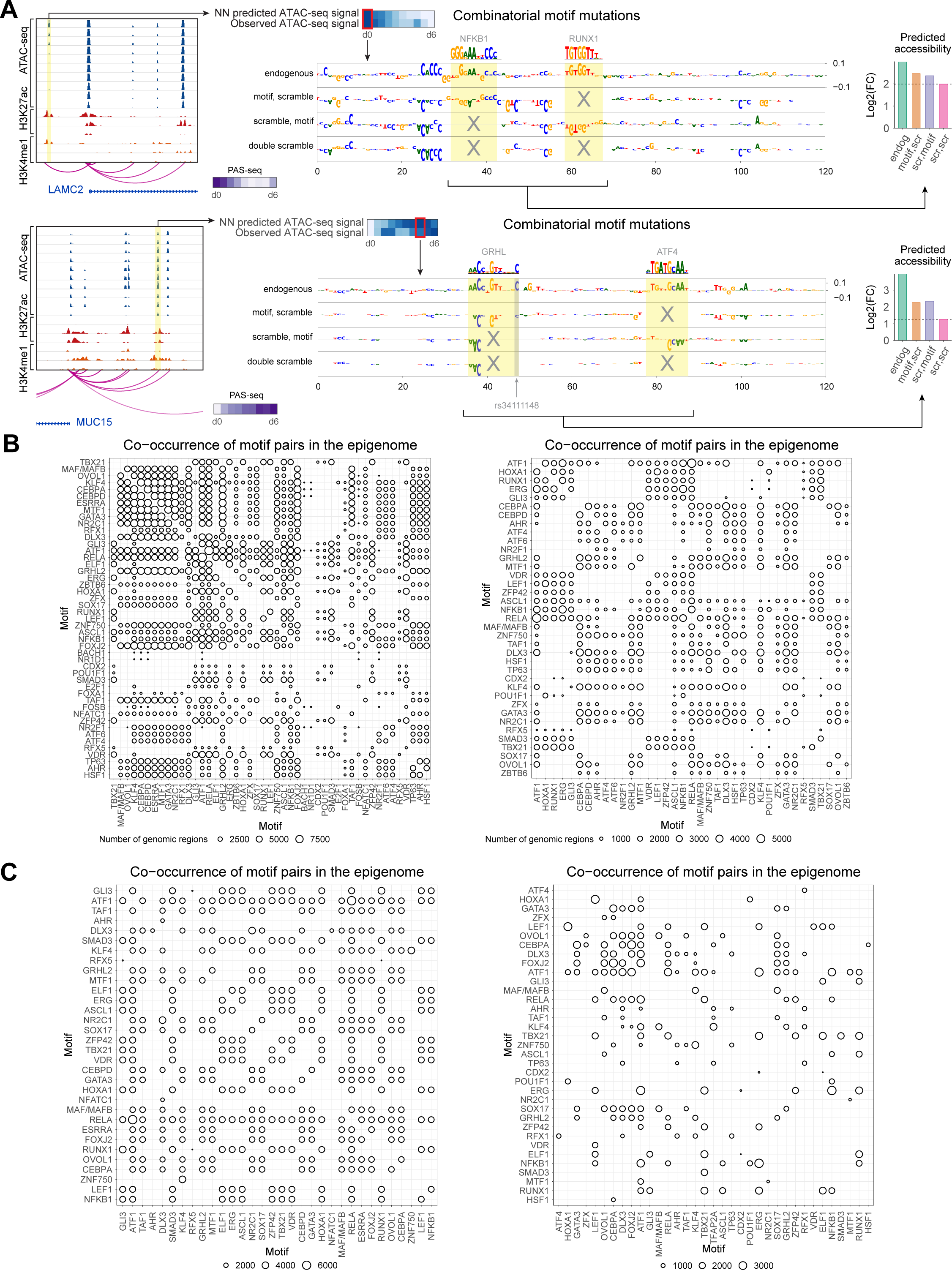
Examples of interacting motif pairs and motif co-occurrence statistics. (**A**) Example regions demonstrating interacting motifs. Top row: putative enhancer affecting LAMC2 gene expression with an interacting NFKB1 motif and RUNX1 motif. The highlighted region in the signal tracks (left) demonstrates correctly predicted ATAC signal by the neural net (top middle heatmap). Base-resolution contribution score tracks are shown for the endogenous sequence and sequences with marginal and joint perturbation of both motifs (middle tracks). The model predicts a super-multiplicative effects of the motif pair on chromatin accessibility (right plot). Bottom row: Analogous plots for a putative enhancer affecting MUC15 gene expression with an interacting GRHL motif and ATF4 motif. (**B**) Co-occurrence statistics (size of circle represents number of instances) for motif pairs based on all motif instances based on sequence matches (left) and motif pairs based on predictive, active motif instances based on contribution score weighted sequence matches (right). Predictive motif instances highlight less promiscuous, more specific co-occurrence statistics. (**C**) Analogous co-occurrence statistics for motif pairs using all motif instances (left) and predictive motif instances (right) after filtering for pairs that show significant GO term enrichments for associated target genes. Once again, more specific co-occurrence patterns are observed for the predictive motif instances.

**Figure S4.**
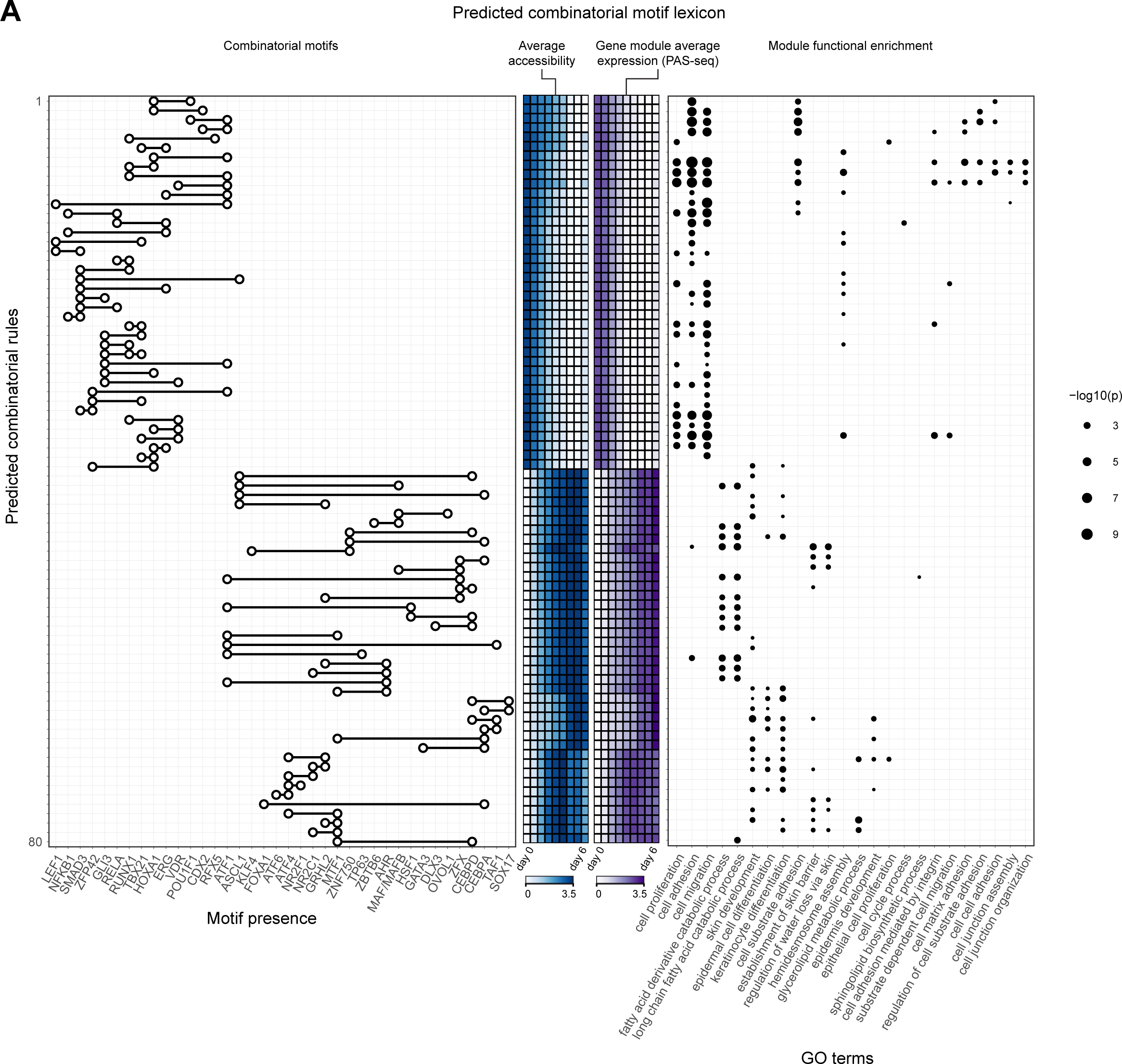
Mapping co-occurring motif pairs to enriched Gene Ontology terms. (**A**) Map of combinatorial rules derived from *in silico* motif interaction analyses. Each row across plots represents a predicted interacting motif pair. From left to right: the motif presence plot demonstrates which motifs are part of the combinatorial rule; the ATAC heatmap demonstrates the average accessibility pattern over CREs containing each motif pair across all time points; the RNA heatmap displays the average gene expression over genes associated with CREs containing each motif pair across the time points; Gene Ontology terms are significantly enriched in the gene sets associated with CREs containing each motif pair.

**Figure S5.**
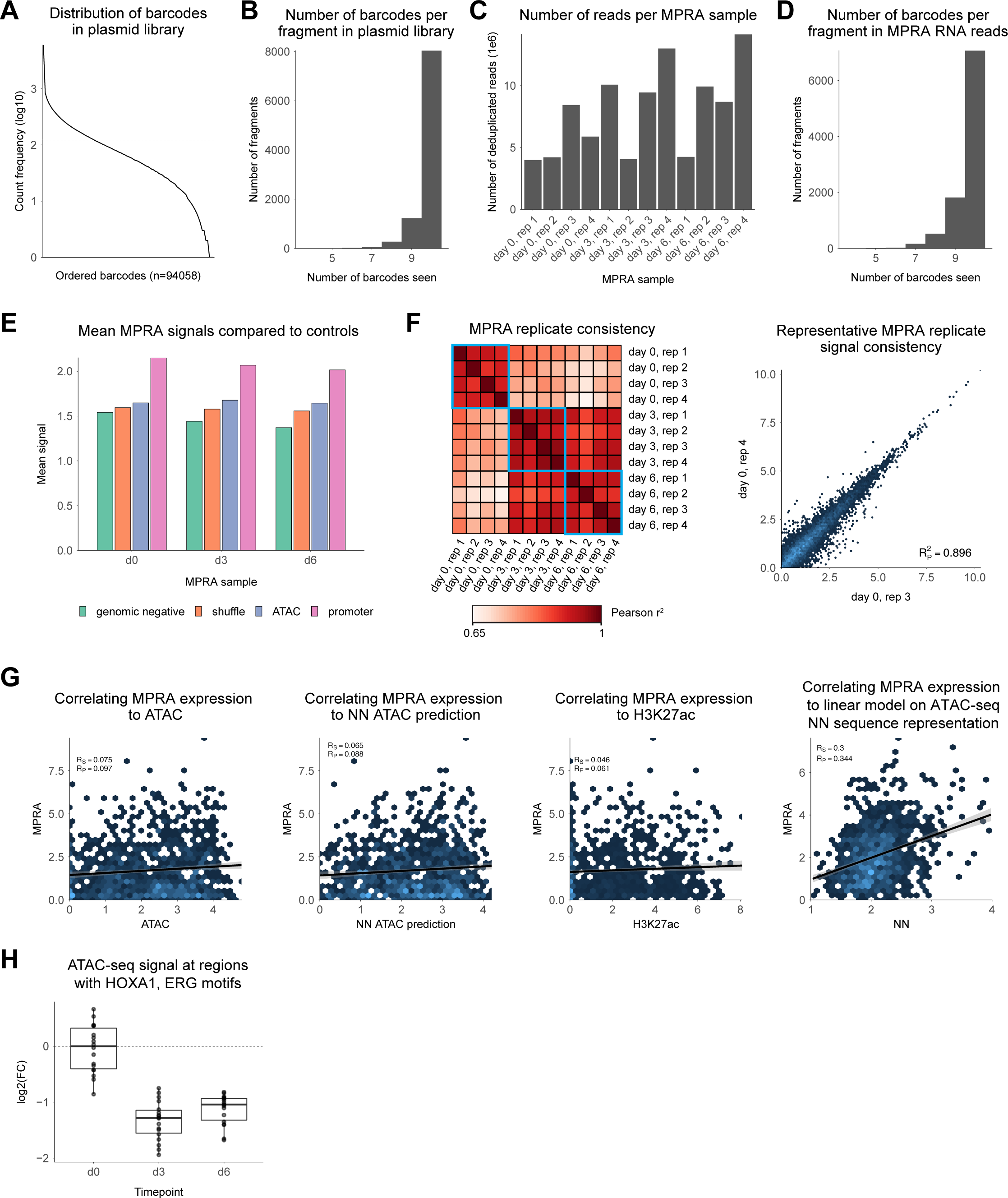
MPRA data quality and comparisons to epigenomic landscapes. (**A**) Distribution of barcodes in plasmid library, demonstrating the skew of barcode representation. (**B**) Number of barcodes per fragment in plasmid library, demonstrating on average 10 barcodes per fragment tested. (**C**) Number of reads per MPRA sample. (**D**) Number of barcodes per fragment in MPRA RNA reads, demonstrating on average 10 barcodes per fragment tested. (**E**) Average MPRA signal compared to controls, showing ATAC regions on average have activity in between negative controls (genomic negatives and shuffled sequences) and positive controls (promoter sequences). (**F**) MPRA replicate consistency. Left: Consistency by Pearson R across replicates and timepoints tested. Right: Consistency of MPRA replicate signal for two example replicates in timepoint day 0. (**G**) Correlation of MPRA signal to various genomic and/or modeling signals: ATAC signal, NN predictions of ATAC signal, H3K27ac, and regression predictions utilizing NN final layer activations as model inputs (results shown on held out test data). (**H**) ATAC signal across timepoints day 0,3, and 6 for sequences containing HOXA1 motif and ERG motif.

**Figure S6.**
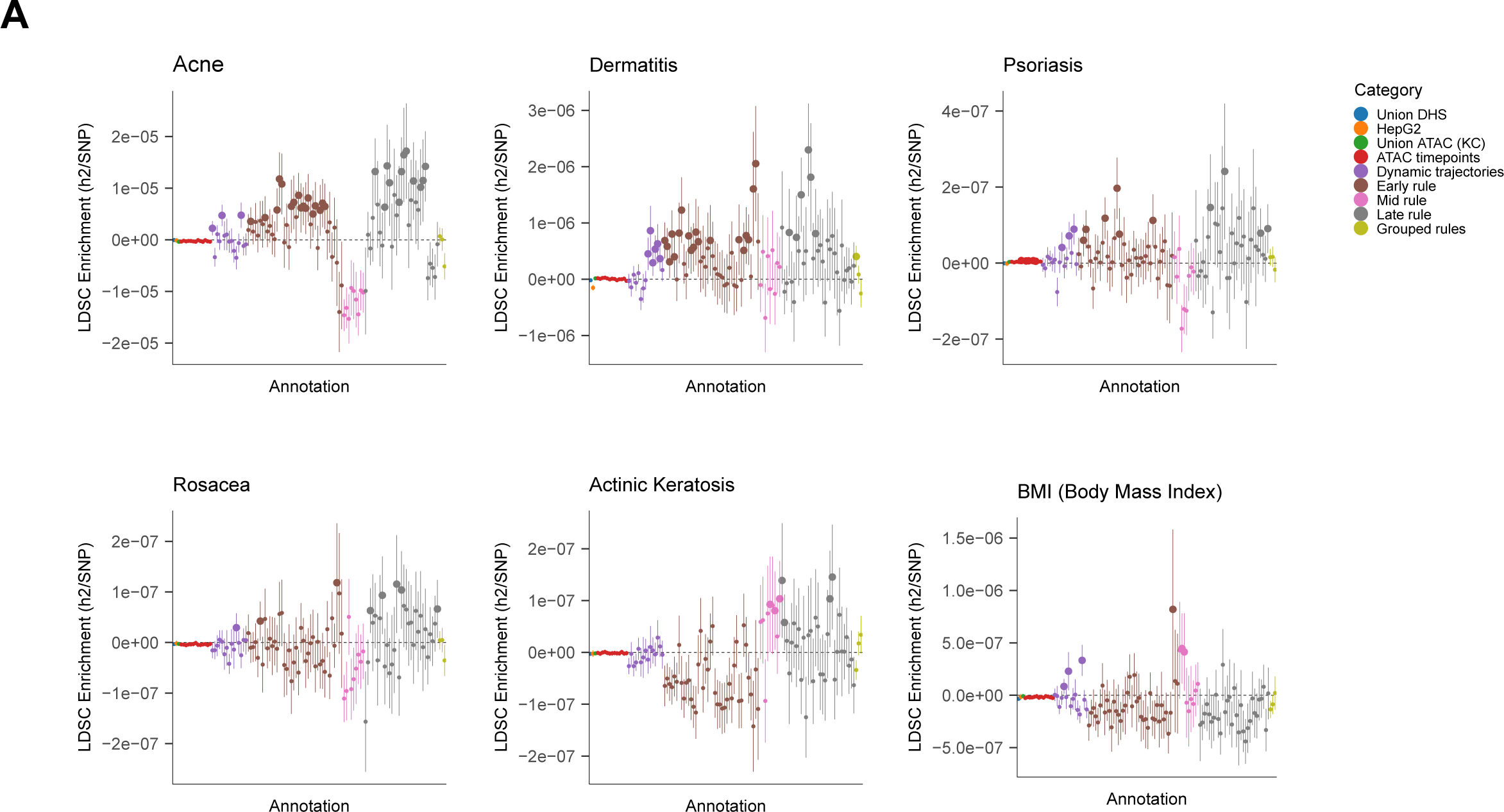
Combinatorial motif pairs are enriched for genetic variants associated with skin phenotypes. (**A**) LD score regression analysis showing differential heritability enrichment of various skin-related diseases and traits in different sets of CRE. The skin phenotypes include: acne, dermatitis, psoriasis, rosacea, actinic keratosis. BMI is shown as a control phenotype. The sets of CREs include: “Early” rules are CREs containing motif pairs that demonstrate decreasing accessibility and activity across the epidermal differentiation time course. “Mid” rules are CREs containing motif pairs those that demonstrate maximal accessibility and activity in the middle of the epidermal differentiation time course. “Late” rules are CREs contained motif pairs that demonstrate maximal accessibility and activity at the end of the epidermal differentiation time course. Union DHS is the union of DNase peaks across all ENCODE DNase datasets. HepG2 are DNase peaks in the HepG2 liver carcinoma cell line. Union ATAC is the union of CREs across all time points of the differentiation time course. ATAC timepoints are the CREs that are accessible in each time point of the epidermal differentiation time course. Dynamic trajectories are clusters of CREs that display specific concordant patterns of dynamic accessibility across the epidermal differentiation time course. Grouped rules are unions of CREs containing motif pairs grouped by different types of dynamic patterns (early, mid, late patterns) across the epidermal differentiation time course.

## SUPPLEMENTARY INFORMATION

## MATERIALS AND METHODS

### Lead Contacts

Further information and requests for resources and reagents should be directed to and will be fulfilled by Lead Contacts Anshul Kundaje and Paul A. Khavari.

### Materials Availability

This study did not generate new unique reagents.

### Data and Code Availability

ATAC-seq, ChIP-seq, and PAS-seq experiments can all be found at the ENCODE portal (https://www.encodeproject.org/). The HiChIP data can be found on at the Gene Expression Omnibus (GEO): GSE158642. The MPRA data can be found on GEO: GSE158477. Integrative analysis code and scripts can be found at https://bitbucket.org/vervacity/ggr-project/, and the deep learning code can be found at https://github.com/kundajelab/tronn.

### Experiments and data processing

#### Cell culture

Primary human keratinocytes were isolated from fresh surgically discarded neonatal foreskin and cultured in Keratinocyte-SFM (Life Technologies 17005-142) and Medium 154 (Life Technologies M-154-500). Pen/Strep (Life Technologies 15140-122) and Anti-mycotic (Life Technologies 15240-062) were also added to the culture. Keratinocytes were induced to differentiate by addition of 1.2 mM calcium (added 12 hours after seeding at confluence) for 6 days in full confluence. Cells were harvested every 12 hours for a total of 13 timepoints and banked into cell pellets, viable batches (10% DMSO in media), or cross-linked with 1% formaldehyde and frozen down at −80 deg C. Further details can be found on the ENCODE portal under GGR experiment accessions (**Table S1**).

#### ATAC-seq experiments

ATAC-seq (Buenrostro et al., 2013) was performed on all 13 timepoints. Detailed methods can be found on the ENCODE portal under GGR experiment accessions (see **Table S1**). ATAC-seq read alignment, quality filtering, duplicate removal, transposase shifting, peak calling, and signal generation were all performed through thes ENCODE ATAC-seq pipeline (https://github.com/ENCODE-DCC/atac-seq-pipeline). Briefly, adapter sequences were trimmed, sequences were mapped to the hg19 reference genome using Bowtie2 (-X2000) (Langmead and Salzberg, 2012), poor quality reads were removed (Li et al., 2009), PCR duplicates were removed (Picard Tools

MarkDuplicates) (2020), chrM reads were removed, reads with MAPQ > 30 were retained and read ends were shifted +4 on the positive strand or −5 on the negative strand to produce a set of filtered high quality reads. These reads were put through MACS2 (Feng et al., 2012) to get peak calls and signal files. Finally, IDR analysis was run on the two replicate peak files to produce an IDR peak file that is the reproducible set of peaks across both replicates (Li et al., 2011).

#### ChIP-seq experiments

ChIP-seq for H3K27ac, H3K4me1, H3K27me3, and CTCF were performed on 3 timepoints (days 0.0, 3.0, and 6.0). Detailed methods can be found on the ENCODE portal under GGR experiment accessions (**Table S1**). ChIP-seq read alignment, quality filtering, duplicate removal, peak calling, and signal generation were all performed through the ENCODE ChIP-seq pipeline. Briefly, sequences were mapped to the hg19 reference genome using BWA (Li and Durbin, 2009), and poor quality reads were removed, PCR duplicates were removed (Picard Tools MarkDuplicates) to produce a set of filtered high quality reads with high mapping scores (MAPQ > 30). These reads were put through MACS2 to get peak calls and signal files. Finally, reproducible sets of peaks across both replicates (naïve overlap peaks) were used for all downstream analysis. The full pipeline can be found at https://github.com/ENCODE-DCC/chip-seq-pipeline2.

#### PAS-seq experiments

PAS-seq was performed on all 13 timepoints. Detailed methods can be found on the ENCODE portal under GGR experimental accessions (**Table S1**). PAS-seq read alignment and quantification were performed using the ENCODE RNA-seq pipeline v2.3.1 (https://github.com/ENCODE-DCC/long-rna-seq-pipeline). Briefly, sequences were mapped to the hg19 reference genome with GENCODE V19 annotations using STAR aligner (v2.4.1d) (Dobin et al., 2013), quantification was performed with RSEM (v1.2.21) (Li and Dewey, 2011), and signal files were produced with STAR and ucsc tools (v3.0.9, http://hgdownload.soe.ucsc.edu/admin/exe).

DESeq2 (Love et al., 2014) was used to identify gene sets that were significantly differentially expressed (adjusted *p*-value < 0.05) in each time point relative to timepoint day 0. We used GSEA (v3.0) (Subramanian et al., 2005) to identify enriched functional terms for each differential gene set. We used the GseaPreranked tool and classic scoring scheme, to determine the GSEA normalized enrichment score (NES) for skin-relevant gene sets from MSigDB (Subramanian et al., 2005), specifically CORNIFIED_ENVELOPE, KERATINIZATION, and KERATINOCYTE_DIFFERENTIATION.

#### HiChIP experiments

The HiChIP protocol was performed as previously described (Mumbach et al., 2016), using antibody H3K27ac (Abcam, ab4729) on 10 million cells per sample with the following modifications. Samples were sheared using a Covaris E220 using the following parameters: Fill Level = 10, Duty Cycle = 5, PIP = 140, Cycles/Burst = 200, Time = 4 minutes and then clarified by centrifugation for 15 minutes at 16100 rcf at 4° C. We used 4 ug of antibody to H3K27ac and captured the chromatin-antibody complex with 34 uL Protein A beads (Thermo Fisher). Qubit quantification following ChIP ranged from 125-150 ng. The amount of Tn5 used and number of PCR cycles performed were based on the post-ChIP Qubit amounts, as previously described (Mumbach et al., 2016). HiChIP samples were size selected by PAGE purification (300-700 bp) for effective paired-end tag mapping and where therefore removed of all primer contamination. All libraries were sequenced on the Illumina HiSeq 4000 instrument to an average read depth of 300 million total reads.

HiChIP paired-end reads were aligned to the hg19 genome using the HiC-Pro pipeline (Servant et al., 2015). Default settings were used to remove duplicate reads, assign reads to MboI restriction fragments, filter for valid interactions, and generate binned interaction matrices. HiC-Pro filtered reads were then processed using hichipper (Lareau and Aryee, 2018) using the {EACH, ALL} settings to call HiChIP peaks to MboI restriction fragments. HiC-Pro valid interaction pairs and hichipper HiChIP peaks were then processed using FitHiChIP (Bhattacharyya et al., 2019) to call significant chromatin contacts using the default settings except for the following: MappSize=500, IntType=3, BINSIZE=5000, QVALUE=0.01, UseP2PBackgrnd=0, Draw=1, TimeProf=1.

### Analysis of epigenomic and transcriptomic landscapes

#### Genome annotations: reference genome, transcription factors, known motifs

We use reference genome hg19 and GENCODE v19 (Frankish et al., 2019). For conversions between Ensemble IDs and HGNC, we use the biomaRt package in R (Durinck et al., 2009). For transcription factors, we use the FANTOM5 list of transcription factors (**Table S9**) (Lizio et al., 2017). For conversion of Entrez IDs to Ensembl IDs, we use the biomaRt package in R.

For our known motif compendium, we use the HOCOMOCO resource (Kulakovskiy et al., 2018). To improve the quality of the motifs, we first remove non-informative bases on the ends of all position weight matrices (PWMs) in the database by clipping positions with information content (IC) < 0.4 from the ends in, until we hit a position with IC > 0.4. We reduce redundancy in this database using the RSAT matrix clustering methodology (Castro-Mondragon et al., 2017). In brief, we cross correlate all motifs to all other motifs in the database, getting both the max raw cross correlation (cor) and the max normalized cross correlation (Ncor). The Ncor is the max cross correlation normalized by a width metric (divide the length of the best cross correlated alignment of the two PWMs by the number of overlapping base pairs between the two PWMs). We use 1 - Ncor as a distance metric to build a hierarchical clustering of the PWMs. We then merge PWMs from the leaves of the hierarchical clustering tree towards the root, stopping at each branch when cutoffs for cor and Ncor (cor < 0.8, Ncor < 0.65) are passed. These cutoffs are the ones empirically derived in the RSAT matrix clustering study. We track all PWMs that were merged as well as associated Ensembl IDs for the corresponding transcription factors.

#### Determining a keratinocyte atlas of cis-regulatory elements

To determine the landscape of accessible regulatory elements across keratinocyte differentiation, we take the union set of the ATAC-seq peaks across all timepoints, using bedtools merge (Quinlan and Hall, 2010), to determine an atlas of cis-regulatory elements (CREs). We use the IDR peak files for each timepoint as the peak set for that timepoint. This CRE atlas consists of 225,996 accessible regions that are accessible at some timepoint in differentiation. At this point in the analysis it was noted that days 3.5, 4.0, and 5.5 had small differences that could be attributed to a growth response from media changes, which were not noted to significantly change the regions included in the CRE atlas but could have important effects on the accessibility signals and downstream quantitative analyses. These timepoints were therefore removed for all downstream analyses. With our valid timepoints we generated a signal coverage matrix with the following computational pipeline. At the biological replicate level, we determined the transposase-corrected cut sites (the single base pair locations of transposase binding events on genomic DNA) from the sequencing reads by taking the read ends and correcting the positions to be +4 on the 5’ end and −5 on the 3’ end. We then count the number of cut sites that fall into each element of our CRE atlas to get transposase events per biological replicate sample. This gives us a count-based matrix of (regions, samples). This count matrix with replicate information can be appropriately analyzed with DESeq2 with its underlying assumptions (Love et al., 2014). We thus use DESeq2 on all pairs of timepoints to get all CREs that have differential signal between any pair of timepoints, using an FDR of 0.0005 to give us a post-analysis Bonferroni corrected FDR of 0.05 across all tests. Under this analysis framework, 47,835 CREs (21% of the CRE atlas) were found to be dynamically accessible across differentiation.

#### Time series clustering of dynamic CREs with replicate reproducibility

To group the dynamically accessible CREs into defined trajectories across time, we utilized Dirichlet Process-Gaussian Process (DP-GP) time series clustering with replicate reproducibility. This analysis framework extends DP-GP time series clustering (McDowell et al., 2018) to consider replicates and to determine which clusters are reproducible across replicates. First, we sum the transposase event counts for each biological replicate into a pooled count for each timepoint. This pooled count matrix is then used to calculate the DESeq2 regularized log transform to get a normalized signal matrix, where each CRE has a normalized value across timepoints. This same regularized log transform is applied to count matrices for replicate 1 and replicate 2, to generate similar normalized signal matrices for each replicate that are all normalized to the same transform. Then, the signal matrix with the pooled data is subsampled (n=5000 for speed, since the algorithm was originally built to run effectively at the scale of thousands of genes, not tens of thousands of regions) with the default parameters, providing the initial set of time series clusters. The cluster set is filtered for cluster size such that any cluster that has a total membership of CREs < 2% of all dynamically accessible CREs is removed. The cluster set is further filtered to remove non-dynamic trajectories, which are the clusters whose multivariate Gaussian process does not reject the null hypothesis of no change across time (in other words, the 0-vector falls in the 99.9% multivariate confidence interval). We then run reproducibility in the following manner. For each CRE, with its corresponding signal trajectory across time, we assign the CRE to each cluster that it could match, and this is also done for the pooled signal trajectory of that CRE as well as the signal trajectories in the separated replicates. The CRE matches a cluster if it’s in the multivariate confidence interval (CI 0.95) for the trajectory, and is correlated by Spearman and Pearson correlation (*p*<0.05). We then only keep cluster matches for that CRE if all three trajectories – pooled, replicate 1, and replicate 2 – were matches in that cluster. If there is more than one matched cluster, the CRE is assigned to the cluster for which it is the least Euclidean distance away from the mean trajectory. If there are no matched cluster, the CRE is considered irreproducible across time and discarded. After this is done for all CREs, any clusters that do not have matched CREs are discarded. This framework thus allows for utilizing replicate information within a time series framework to improve clustering as well as cluster membership. Under this analysis, 15 time series patterns of accessibility were found in keratinocyte differentiation, comprising 40,103 dynamically accessible and time series reproducible CREs.

#### Analysis of histone modifications in the CRE atlas

To characterize the diversity of CREs by histone modification, histone marks were analyzed with an accessibility-centric approach. For each histone mark (H3K27ac, H3K4me1, and H3K27me3), a union set of regions was generated by taking the CREs, extending the flanks on either side by 1kbp, and keeping any CREs that overlapped peaks for that mark across any of the timepoints. This analysis finds 83,785 CREs marked by H3K27ac, 122,395 CREs marked by H3K4me1, and 36,084 CREs marked by H3K27me3. We then generated count matrices for each set of CREs in the following manner. At the biological replicate level, we determined the midpoints from the paired sequencing reads as estimated positions where the histone was present on genomic DNA. We then count the number of read midpoints that fall into each flank-extended element of each CRE to get marked histone events per biological replicate sample. This gives us a count-based matrix of (regions, samples). This count matrix with replicate information can be appropriately analyzed with DESeq2 with its underlying assumptions (Love et al., 2014). We thus use DESeq2 on sequential pairs of timepoints to get differentially marked CREs across time. Given three possible transitions (increase, no change, decrease in histone mark signal), and three timepoints (day 0, 3, and 6) for which histone mark data was collected, we enumerate 9 possible patterns for histone marks across time.

#### Analysis of chromatin states in the CRE atlas

To characterize the diversity of CREs by chromatin state, the histone mark analysis from above was used to consider all histone marks together. Chromatin states were generated by enumeration. With 9 possible patterns for each histone mark and three assayed marks, the total possible chromatin states is 729. However, most of the possible states do not appear, demonstrating a much more limited set of states.

#### Determining the transcriptomic atlas of keratinocyte differentiation

To determine the landscape of transcripts across keratinocyte differentiation, we first determine the set of expressed genes at each timepoint. We do this by first normalizing the full matrix of protein-coding transcripts across timepoints using the rlog function from DESeq2 (Love et al., 2014), and then setting an empirical threshold based on the best separation of a Gaussian mixture model on the rlog normalized values (threshold = 4.0). We then take the union of all expressed genes across timepoints to determine the transcriptomic atlas, which consists of 12,190 genes. We then use DESeq2 on all pairs of timepoints to get all genes that have differential signal between any pair of timepoints, using an FDR of 0.0005 to give us a post-analysis Bonferroni corrected FDR of 0.05 across all tests. Under this analysis framework, 5,046 genes (41% of the transcriptome atlas) were found to be dynamically accessible across differentiation.

#### Time series clustering of dynamic genes with replicate reproducibility

To group the dynamic genes into defined trajectories across time, the same framework used for the dynamic CREs was also utilized for the dynamic genes (see above section, “Time series clustering of dynamic CREs with replicate reproducibility”). Under this analysis, 11 time series patterns of expression were found in keratinocyte differentiation, comprising 3,610 genes (29% of the transcriptomic atlas) that are dynamic and time-series reproducible.

#### Analysis of chromatin conformation

To determine a set of loops for downstream analyses, a replicate-based analysis was run to get replicate reproducible loops. For each timepoint, loops were generated for the pooled data (aggregated across both replicates), replicate 1, and replicate 2. Consensus loops were generated by getting the consensus endpoints from the union merge across the pooled, replicate 1, and replicate 2 endpoints. These were filtered such that each loop had a non-zero value for the pooled version, replicate 1 version, and replicate 2 version. These values were run through IDR (*p*<0.05) to keep loops that were replicate consistent (Li et al., 2011). These loops were then merged across timepoints to get the union set of replicate consistent loops across differentiation. Under this analysis, 101,884 loops were replicate consistent.

#### Linking by proximity

We utilize an exponential decay function e(−*λd*) to compute a linkage score between each ATAC-seq peaks and each expressed genes separated by distance *d*. Since previous work has shown that the median distance for functional distal regulatory elements to gene TSSs is 25kb (Gasperini et al., 2019), we fit the exponential decay function such that the median score is at 25kb (i.e., *λ* = ln(2)/25000). We then keep all peak-gene links that are within 100kb of each other. For curating a gene set linked to a region set, we use the above links to get genes that are proximally linked to the regions where 1) the genes are expressed at some point in the timecourse, 2) the gene TSS is within 100kb upstream or downstream of a region, 3) the summed score for the gene is > 0.5. For example, if two regions are within 100kb of a gene TSS and the sum of the link scores for the two regions is 0.51, then the gene is kept as part of the downstream gene set, with the corresponding summed score. We then use this summed score to rank the genes so that a ranked enrichment tools can be used. This linking and scoring strategy is the main strategy used to find gene sets and gene set enrichments.

### Deep learning on dynamic regulatory DNA sequence

#### Convolutional neural networks on DNA sequence

We trained multi-task convolutional neural networks (CNNs) to accurately map 1 kbp DNA sequence regions across the genome to quantitative read outs of chromatin accessibility and multiple histone marks in each time point of keratinocyte differentiation. CNNs can learn complex sequence patterns that are predictive of genome-wide chromatin accessibility and histone mark profiles. We use a multi-stage, transfer learning training regimen to maximize prediction performance and model stability by leveraging large compendia of chromatin accessibility data across 100s of diverse tissues.

#### Architecture

We used the previously optimized multi-task Basset CNN architecture for predicting genome-wide chromatin accessibility from DNA sequence across multiple samples (Kelley et al., 2016). The inputs to the model are 1 kbp long DNA sequences that are one-hot encoded (A=[1,0,0,0], C=[0,1,0,0], G=[0,0,1,0], T=[0,0,0,1]). The Basset model has three convolutional layers with the following parameters: the first layer has 300 filters of size (1, 19) and stride (1, 1) followed by batch normalization, a ReLU non-linearity, and max-pooling with size (1, 3) and stride (1, 3); the second layer has 200 filters of size (1, 11) and stride (1, 1) followed by batch normalization, a ReLU non-linearity, and max-pooling with size (1, 4) and stride (1, 4); the third layer has 200 filters of size (1, 7) and stride (1, 1) followed by batch normalization, a ReLU non-linearity, and max pooling with size (1, 4) and stride (1, 4). After the convolutional layers there are two fully connected layers, each with 1000 neurons, followed by batch normalization, a ReLU non-linearity, and dropout where the keep probability is 0.7. The final layer mapped to multiple outputs (multi-task output) spanning the time points and each of the different types of molecular read outs (chromatin accessibility or histone marks). We use binary or continuous output labels and associated loss functions in the multi-stage training (see below). When training on binary labels (accessible vs. not accessible or bound vs. unbound), we use the binary cross-entropy loss function with logistic outputs. When training on continuous, quantitative measures of accessibility or histone marks, we use the mean-squared error loss function with linear outputs. The multi-task loss is the sum of the loss over all tasks.

#### Multi-stage transfer learning regimen

We bin the genome into 1 kbp windows with a stride of 50 bp. Each bin can serve as an example in a training, validation/tuning or test set. We divide chromosomes into 10 folds (**Table S6**). We use a cross-validation set up where we use 8 folds for training, 1 for validation/tuning, 1 for testing.

We use a multi-stage training regimen to maximize performance and model stability. In stage 1, we train a ‘reference’ multi-task CNN model with randomly initialized parameters (variance scaling initialization, ie Xavier intialization) on DNase-seq and TF ChIP-seq data from a large collection of biosamples from the ENCODE and Roadmap Epigenomics Project (ENCODE Project Consortium, 2012; Kundaje et al., 2015). All datasets used are detailed in **Tables S7, S8**. In this stage, the labels associated with each input sequence are binary. A 1 kbp sequence in the genome is assigned a positive label for a particular task (DNase-seq or TF ChIP-seq in a specific biosample), if the central 200bp of the sequence overlaps a DNase-seq or TF ChIP-seq peak in the biosample by at least 50%. All other bins in the genome are assigned negative labels for that task. The possible negative labeled bins significantly outnumber the positive labeled bins, since much of the genome is not accessible (or not TF bound). Hence, we use a subset of informative negative examples from the training chromosomes to train the models. For each task, we include negative labeled bins flanking every positive labeled bin (3 flanks, stride 50bp, on either side of the region). We further sample negatively labeled bins (half as many positive bins in the task). Finally, we include bins that overlap a comprehensive catalog of DNase-seq peaks (Kundaje et al., 2015). This generates a dataset with reasonable class imbalances per task while maintaining diversity in negative examples (Kim and Kundaje, 2020a).

In stage 2, we initialize a multi-task CNN model with the parameters derived from the reference ENCODE/Roadmap model (Kim and Kundaje, 2020b) and then train it to map DNA sequence bins to binary labels corresponding to important region sets as derived in the characterization of the epigenomic landscape. These important region sets include: ATAC-seq, H3K27ac, H3K4me1, and H3K27me3 region sets by timepoint, the region sets defined by accessibility time series clustering, region sets defined by dynamic and static histone modifications, region sets defined by dynamic and static chromatin states, and region sets from TF ChIP-seq experiments for CTCF, TP63, ZNF750, POL2, and KLF4. In total these region sets comprise 119 binary label sets used for multitask training. The genomic bins for training and their associated binary labels for each of the tasks are constructed as described above (Kim and Kundaje, 2020c).

In the final stage 3, we initialize a multi-task CNN with the parameters of the binary keratinocyte model from stage 2 (Kim and Kundaje, 2020d). We then train the model CNN with the mean-squared error regression loss function to map DNA sequence bins to continuous, quantitative measures of ATAC-seq, H3K27ac ChIP-seq and H3K4me1 ChIP-seq in our keratinocyte differentiation time course (19 tasks). The genomic bins used for training cover the union of peaks across all time points. For each 1 kbp sequence bin, we compute the average of the log of the smoothed depth-normalized read coverage (log of the MACS2 fold-enrichment of smoothed observed 5’ end counts relative to expected local Poisson background) over the central 200 bp of the bin for ATAC-seq or over the entire 1 kbp for histone marks. The average is computed using bigWigAverageOverBed (column mean0). The average signal scores are normalized using quantile normalization across all time points for each of the assays (ATAC, H3K27ac, and H3K4me1). These normalized scores are used as quantitative labels for each bin.

We use the same cross-validation folds for training, tuning and testing across all stages. The model parameters are transferred across stages to exactly match the cross-validation fold structure. Hence, for each fold, the test sets are completely held-out across all stages of training. This multi-stage training set up allows the model to utilize larger sets of existing data to improve its understanding of DNA sequence and regulatory logic encoded in the human genome.

#### Training hyperparameters

The following hyperparameters were used for all models at all stages. We train for a maximum of 30 epochs with early stopping, where the patience (number of epochs of nonimproved performance before stopping) is 3 and the metric considered is average AUPRC across all tasks on the validation set. The loss function for classification models is binary cross entropy, and the loss function for regression is mean squared error (MSE). The optimizer used is RMSprop with a learning rate of 0.002, a decay of 0.98, and a momentum of 0.0.

#### Performance evaluation

We evaluate on the held-out test chromosomes of each fold, calculating our performance metrics across the entire length of the chromosomes (genome-wide evaluation). For each task, we use the area under the precision-recall curve (AUPRC) to measure performance of the binary models and Spearman’s R and Pearson’s R to measure performance of the regression models.

#### Prediction calibration through quantile normalization

Using MSE loss on regression models provides effective ranking across predictions in the same task, but the prediction outputs may not be well calibrated to match the observed output labels. As such, we rescale the model’s continuous output predictions by quantile normalizing the distribution of the model predictions with respect to the distribution of the ground truth measured labels. We obtain prediction scores for a random set of 1000 examples, which then provides us with a distribution of predicted scores and a corresponding distribution of the labels. We can then use those distributions to quantile match a prediction score value to a label score (for example, we can determine that a prediction score is in the 90th percentile of the distribution of prediction scores and should be matched to the 90th percentile of the distribution of label scores). Importantly this re-scaling does not actually change the performance of the model, it simply re-calibrates the output. Additionally, given that the continuous signal labels across tasks are quantile normalized relative to each other, the re-calibration of the prediction scores also normalizes the prediction scores across tasks.

### Inference of predictive motif instances

#### Overview

The multi-task CNNs, described above, map every candidate regulatory DNA sequence to quantitative measures of chromatin accessibility at each time point in the differentiation time course. We developed an interpretation framework to interrogate the model and decipher motif instances in each candidate element that are predictive of chromatin accessibility at each time point. First, we use gradient based feature attribution methods to decompose the predicted output (at each time point) for an input sequence in terms of contribution scores of each nucleotide in the sequence. We develop methods to stabilize and normalize the scores. We develop stringent null models to identify statistically significant contribution scores. We then use a large compendium of pre-compiled TF motifs to scan and score the sequences as well as the contribution score profiles. We develop stringent null models to infer predictive motif instances that have statistically significant contribution scores and sequence match scores. The following sections provide details for each of these steps.

#### Estimating nucleotide-resolution contribution scores

The gradient of the predicted output with respect to each base at each position in the input DNA sequence, gated by the observed base, estimates the sensitivity of the output to infinitesimal changes in the input (Simonyan et al., 2014). This measure of importance is often referred to as input-gated gradients. The method is efficient since a single backpropagation pass can be used to estimate the contribution of all nucleotides in an input DNA sequence to a specific output prediction.

We compared the input-gated gradient scores to contribution scores derived from another related approach called DeepLIFT (Shrikumar et al., 2017) on a subset of the time points. DeepLIFT backpropagates a score, analogous to gradients, which is based on comparing the activations of all the neurons in the network for the input sequence to those obtained from neutral ‘reference’ sequences. We use 12 dinucleotide-shuffled versions of each input sequence as reference sequences. We used the DeepSHAP implementation of DeepLIFT (https://github.com/slundberg/shap/blob/0.28.5/shap/explainers/deep/deep_tf.py) to obtain contribution scores for all observed bases in each sequence.

We estimated input-gated gradient and DeepLIFT contribution scores for all nucleotides in all sequences with respect to quantitative chromatin accessibility predictions for three time points (0h-early, 3h-mid and 6h-late), using each of the models for the 10 folds of cross-validation. For each method, we averaged the scores for each sequence in each time point across all the 10 folds. For each sequence, we used cosine similarity to compare the average input-gated gradient and DeepLIFT score profiles separately. We observed high similarity between input-gated gradients and DeepLIFT scores (median cosine similarity across all sequences and all the 3 time points = 0.8736). While gradient based scores are often more unstable and less accurate than DeepLIFT scores, the regularization of our models via the multi-stage transfer learning and averaging over folds, greatly stabilizes the gradient based scores. Hence, we decided to use input-gated gradient scores as contribution scores for all downstream analyses, since it is more efficient than DeepLIFT and produces very similar contribution score profiles with respect to motif instance discovery.

#### Estimating statistically significant contribution scores

For each input sequence, we compute input-gated gradient score profiles from dinucleotide shuffled versions of the sequence. We use these scores to construct an empirical null distribution of contribution scores for that sequence. We use that empirical null distribution to derive empirical statistical significance of the observed contribution scores. We use a threshold of *p* < 0.01 to call statistically significant scores. The scores of all positions that do not pass the significance threshold are set to 0.

#### Normalization of contribution scores

We normalize the contribution score profile of each sequence by dividing the score of each position by the sum of the absolute value of contribution scores across the entire sequence and multiplying them by the predicted output.

#### Trimming contribution scores

We observed that statistically significant contribution scores peaked within 160 bp for the peak summit. Hence, we trim the DNA sequences from its original 1000bp context to the central 160bp for downstream analyses. We further eliminate the trimmed sequences from downstream analyses that have less than 10 base pairs of significant scores. These shorter sequences are also compatible with testing in reporter constructs.

#### Average contribution score profiles across all folds for each sequence in each time points

We estimate contribution score profiles for each sequence with respect to predictions in each of the time points using models from each of the 10 folds. These score profiles are filtered for statistical significance, normalized and trimmed as described above. We average the contribution scores of each position in each sequence for each time point, across the 10 folds. We compute the 99% confidence interval for each position using the scores from the 10 folds. If the confidence interval includes 0, the score of the position is set to 0, else it is set to be the average score across the 10 folds.

#### Validation of contribution scores by Allele-sensitive ATAC (asATAC) analysis

We utilized our ATAC-seq data to determine allele-sensitive accessible sites. Since the data was collected from primary samples, we were able to utilize an allele-sensitive ATAC-seq analysis to determine a number of single nucleotide polymorphisms (SNPs) that exhibited significant allelic imbalance of ATAC-seq reads. Utilizing QuaSAR (Harvey et al., 2015), a computational framework for calling genotypes and allelic imbalanced sites, we were able to call 16,686 heterozygous SNPs and capture 283 SNPs with statistically significant (FDR < 10%) allelic sensitivity across our two patient samples (and across all ATAC timepoints). We estimated contribution score profiles for the sequences containing each of these 16,686 SNPs, to determine if the SNP locations overlapped statistically significant contribution scores and whether the models predicted differential accessibility prediction for the two alleles (**Fig. S2D**).

#### Identifying dynamic predictive motif instances using sequence match and contribution scores

We identify dynamic predictive motif instances in each input sequence across time points, for each of the known motifs in the motif compendium, by scanning and scoring the sequence as well as the dynamic the contribution score profiles derived from the model.

First, for each PWM motif, we compute sequence match scores at every position in each sequence. The scanning and scoring can be implemented as a convolution operation. Hence, we use the deep learning framework to implement a single convolutional layer with filters corresponding to each of the PWMs in the deep learning framework. When loading the PWM weights into the filters, we pad the weights to get all filters to be the same size, normalize by the length of the nonzero weights (divide by the length), and convert the weights to a unit vector (divide by L1 norm). We use the convolutional layer to scan and score all PWMs across the forward and reverse complement of each one-hot encoded sequence. We also use the same operation to scan and score dinucleotide shuffled versions of each of the genomic sequences. We thus obtain an empirical null distribution of match scores for each PWM for each sequence. We identify positions with significant sequence match scores as those that pass *p* < 0.05 based on the empirical distributions. For any sequence, the significant positions based on sequence match scores will be identical across all time points.

Next, we use the PWMs to scan and score the dynamic contribution score profiles for each sequence in each time point. Essentially, we repeat the same convolution operation using PWM filters but using the contribution score profiles to weight the one-hot encoded sequences and their reverse complements. Hence, we obtain contribution weighted match scores to the PWMs. We once again retain statistically significant contributed weighted match scores, using a p < 0.05 threshold, based on null distributions of the contribution weighted match scores for dinucleotide shuffled sequences. We compute these contribution weighted match scores and significance separately for the positive contribution scores and negative contribution scores, as negative scores can influence PWM weights detrimentally (e.g. negative PWM weight value that now contributes positively because of a negative contribution score.)

Our final set of predictive motif instances for each sequence in each time point correspond to positions that have significant sequence match scores and significant contribution weighted match scores. Since the contribution score profiles for each sequence can change across time points, the predictive motif instances are dynamic across time points.

#### Identifying significant differential motifs between two sets of sequences

We developed an approach to identify significant differential motifs between any foreground set of sequences relative to a background set of sequences. We use the sequences belonging to each dynamic trajectory as foreground region sets. First, we identify predictive motif instances for all PWMs in both sets using the method described above. We then use bootstraps of GC-content matched background genomic sequences (n=1000) that do not overlap any accessible peaks to estimate a null distribution of the number of PWM hits (average across the sequences in the bootstrapped background set). We then estimate an empirical *p*-value for each PWM in the foreground relative to these bootstrapped backgrounds. We use Storey’s *q*-value method to perform a multiple hypothesis correction. We use an *q*-value threshold of 10% to identify statistically significant differential PWMs.

#### Validation of predictive motif sites by TF ChIP-seq

The predictive motif instances are a subset of all sequence-based motif instances that also have significant contribution scores. We hypothesized that predictive motif instances are more likely to distinguish those that are bound by TFs from unbound motif instances. Hence, we used publicly available TF ChIP-seq to analyze occupancy over these motif sites. First, for each TF with available ChIP-seq data, we used our models to obtain all predictive motif instances of the PWM for the TF. We also collated a control set of motif instances with significant sequence match scores that are not marked as predictive and are matched for accessibility to the set of predictive instances. We matched for accessibility to account for confounding effects of differentially accessible regions. Using pre-processed normalized (MACS2 derived fold enrichment of smoothed 5’-end read coverage relative to local Poisson background) bigwigs from Cistrome (Liu et al., 2011), we contrasted the average the ChIP-seq signal profile over predictive motif instances versus control motif instances using a +/-1 kbp window (20 p bins) around the instances (Ramírez et al., 2016).

#### Validation of predictive motif sites by ATAC-seq footprinting analysis

As above, we separated motif instances of each PWM into two sets depending on whether they were marked as predictive or not, matched for GC content of the surround sequencing context. We then utilized the HINT footprinting tool to generate average ATAC-seq bias-corrected cut site coverage profiles over the two sets of motif instances (Li et al., 2019) using a +/-250 bp window around the motif. We normalized the average footprint profile by computing the fold enrichment of average footprint signal at each position relative to a reference. The reference was computed by averaging the footprint signal in the 50bp flanks on either side of the 250 bp footprint window. We computed the ‘average footprint height’ as the area under the normalized footprint profile in a +/-100 bp window around the motif center, excluding the central +/15 bp around the motif center. We computed the ‘average footprint depth’ as the area under the local maximum of the normalized average footprint profile within +/-10bps on either side of the motif center.

#### Identifying putative TFs binding the motifs based on correlation of weighted PWM scores and TF expression

Since the contribution score profiles for each sequence are dynamic across the time points, the predictive motif instances within each sequence also have dynamic contribution scores across the time points. For each PWM, we identify the locations of all predictive motif instances across all the time points. Each instance is represented by an instance activity vector across time consisting of contribution scores (sum over all positions in the instance) for each of the time points. For a pre-defined set of sequences, we first identify all significant differential motifs relative to a background set (as described above). For each motif, we obtain a motif activity vector for the set of sequences as the average of the activity vectors over all its predictive instances in those sequences. We identify all candidate TFs associated with the motifs (TFs of the same family often bind similar motifs). For each candidate TF, we extract the RNA-seq expression profile (variance stabilized rlog transformed counts from DESeq2) across the time points. We compute the Pearson correlation between the TF expression profile and its motif activity vector. We retain TFs that are expressed in keratinocytes (> 1 TPM) and exhibit a correlation of at least 0.75 with the activity vectors of associated motifs.

### Estimating interaction effects between motifs

### *In silico* mutagenesis scores for motif instances

We use each set of candidate regulatory elements that belong to each trajectory of accessibility dynamics as a foreground set of sequences to identify significant differential motifs relative to the background set of all peaks. As described above, we identify predictive motif instances of differential motifs by scanning sequence and gradient based contribution score profiles with the known motif compendium. We use an *in-silico* motif mutagenesis approach to further corroborate and filter high-confidence predictive motif instances. Specifically, we expect *in-silico* perturbation of predictive motif instances to induce (1) a significant change in the predicted output of the model,(2) a significant change in the contribution scores across the sequence.

We use two complementary approaches to perturb motif instances. The first approach called the “motif scramble” method, randomly scrambles the 10 bp sequence around the center of a predictive motif instance. The scramble maintains the sequence composition of the window while destroying the precise sequence of the motif instance. The second approach, called the “point mutation” method, mutates the most influential base pair with the highest contribution score in the 10bp window around the center of the motif instance. The position is mutated to the base with the most detrimental predicted effect i.e. the base with the most negative gradient score at that position. For both types of mutations, we compute the ‘mutagenesis effect size of a motif instance’ as the difference in the predicted output (units of log depth normalized coverage of ATAC-seq signal) of the model for the mutated sequence relative to the wild type. We also recompute the contribution score profile of the mutated sequence and record the difference in contribution score (‘delta contribution score’) of each position in the sequence relative to the wild type contribution score profile.

We generate null control distributions for the mutagenesis effect sizes and change in contribution scores as follows. First, we identify 10 randomly chosen positions that have non-significant contribution scores (set to 0 after thresholding for significance) in the central 200 bp of each wild type sequence. We expect mutations to these positions to have no significant effect on the output or contribution scores of other positions in the sequence. We mutate (point mutation or scramble around) each of these 10 positions and compute the difference in the output prediction as well the delta contribution scores for all the other positions in the sequence. We fit separate Gaussian null distributions to the mutagenesis scores and the delta contribution scores from these 10 expected null mutations. The use these null distributions to estimate the statistical significance (*p*-value < 0.1) of mutagenesis scores and delta contribution score profiles for each predictive motif instances.

We identify candidate epistatic partners of a motif instance in a sequence, as all other predictive motif instances in the sequence that overlap positions with significant delta contribution scores of a target motif instances in the sequence. We have previously described this approach as Deep Feature Interaction Maps (Greenside et al., 2018).

#### Functional enrichment of co-occurring pairs of predictive motifs

We associate each of the regulatory sequences of accessible peaks supporting a combinatorial motif set to proximal genes as follows. For sequence, we first identify up to two closest candidate genes (based on distance from TSS) within a +/-500 kb of the sequence, such that the genes are expressed in at least one of the time points in our differentiation time course. We then restrict all peak-gene associations to those that exhibit significant correlation between the ATAC-seq enrichment of the peaks (log fold enrichment) and the RNA-seq (TPM) expression levels of the genes across the time course. We thus obtain a gene set that is putatively regulated by any combinatorial motif set. We then test the gene sets for enrichment of functional annotations using gProfiler (Reimand et al., 2016). We use a background set of all genes expressed at any timepoint in differentiation time course) to get functional enrichments. We keep combinatorial rules that are functionally enriched for skin-related terms. We also combine combinatorial rules that were discovered in different trajectories but marked with the motif combination to create a set of rules that are all distinct motif combinations.

#### Testing interaction effects between pairs of motifs with combinatorial *in silico* mutagenesis

Co-occurring pairs of predictive motifs in a regulatory sequence can have different types of quantitative joint effects on chromatin accessibility (depth normalized ATAC-seq read coverage). We explore three types of joint effects. Lack of motif interactions would manifest as independent, additive effects on coverage. Interactions between motifs learned by the model would manifest as multiplicative (additive in log space) or super-multiplicative effects (multiplicative in log space) on coverage. For all pairs of functionally enriched pairs of co-occurring motifs, we identified all the sequences containing predictive instances of the pair (as described above). We then used two complementary approaches to test each instance of a pair of motifs for epistatic interactions.

First, we used the Deep Feature Interaction Map method (Greenside et al., 2018) to score epistatic interactions between pairs of candidate predictive motif instances (say A and B) in a sequence. Specifically, as described in the motif ISM section, we infer the positions in the sequence that exhibit statistically significant delta contribution scores due to *in silico* mutations to motif A. If motif instance B overlaps any positions with significant delta contribution scores then it is estimated to have an interaction effect with motif A on ATAC-seq read coverage.

Next, we corroborate the DFIM scores, with an explicit combinatorial *in silico* motif mutagenesis approach using both the ‘scramble’ and ‘point mutation’ approach. Assume we have two motif instance A and B in a sequence that we would like to test for epistatic interactions using the model.

- We record the model’s output with both motif instances intact in the sequence =*o*.
- We record the output after ‘mutating’ only motif A i.e. the sequence only contains an intact motif B = *b*.
- We record the output after mutating only motif B, i.e. the sequence contains an intact motif A = *a*.
- Finally, we record the output after mutating both motifs A and B, which is a baseline = *n*.
- We compute the marginal effect size of adding motif A relative to a null sequence that does not contain either of the motifs = *(a – n)*.
- We compute the marginal effect size of adding motif B relative to a null sequence that does not contain either of the motifs = *(b – n)*.
- We compute the joint effect of adding motif A and B relative to the sequence that does not contain either of the motifs = *(o – n)*

We then compare the joint effect size *(o – n)* to the sum of the marginal effect sizes *(a – n) + (b – n) = (a + b – 2n)*. We run a Wilcoxon signed-rank test on the paired values (joint vs. sum of marginals) across all instances of a motif pair to determine whether the joint effects on the motif pair instances is significantly greater or less than the sum of the marginal effects.

Since the output predictions are in units of log depth normalized coverage, additivity in log units translates to multiplicative effects in units of coverage. If the joint effect is significantly larger than the sum of the marginal effects, motifs A and B have super-multiplicative effect on coverage. If the joint effect is significantly lower than the sum of the joint effects, motifs A and B exhibit a sub-multiplicative effect on coverage. A non-significant difference between the joint and sum of marginals indicates a multiplicative effect of motif A and B on coverage.

### Design of Massively Parallel Reporter Assay (MPRA) to test intrinsic activity dynamics of combinatorial motif rules

#### MPRA design

We designed MPRA constructs guided by the combinatorial motif sets that have positive motif interaction scores using the ‘motif scramble’ and ‘point mutation’ motif perturbations. For each rule of interacting motif pairs, we randomly select 19 genomic subsequences of length 160 bp within accessible peaks, contain predictive instances of both motifs in the rule and exhibit positive interaction scores. We test the endogenous sequence and all versions of the sequences in which the motifs are combinatorially mutated.

This sampling design allows us to test the following hypotheses:

1. Trajectory: does the motif combination produce a reporter activation pattern across time points (days 0, 3, and 6 in the in vitro model) that was predicted by the trajectory it was derived from?
2. Interactions: do the motif pairs exhibit multiplicative or super-multiplicative interaction effects on intrinsic reporter activity?

We include the following positive and negative controls. As positive controls, we use 316 TSSs of the highest expressed genes (at any time point in skin differentiation). As negative controls, we generate dinucleotide shuffled versions of 50 randomly selected genomic test sequences selected above. We also select 50 negative controls from the genome that are not found in the master list of accessible regions across keratinocyte differentiation. The list of all constructs are in **Table S10**.

#### Library cloning

The MPRA oligo library was synthesized using Agilent’s oligo library synthesis platform. Each oligo sequence consisted of: 5’-FWD primer binding site-[ACTGGCCGCTTCACTG]-176 nt insert-XhoI-NheI-20 nt oligo barcode-REV primer binding site-[AGATCGGAAGAGCGTCG]-3’. The oligo library was amplified using the FWD primer 5’-GCTAAGGAATTCACTGGCCGCTTCACTG-3’ and REV primer 5’-GCTAAGGGATCCCGACGCTCTTCCGATC-3’, which add EcoRI and BamHI restriction sites, respectively. The resulting PCR product was gel purified, digested with EcoRI and BamHI, and ligated into the pGreenFire1 lentivector backbone. Takara Stellar competent cells were transformed with the plasmid library and a fraction of the bacteria were plated to ensure a library coverage of at least ten-fold. The remainder of the transformation was incubated overnight in Luria broth. Plasmids were isolated using the Qiagen Plasmid Plus Maxi kit. If insufficient colonies were obtained to ensure a library coverage of at least ten-fold, additional transformations were performed and plasmid preps were pooled. In the second cloning step, the plasmid library was digested with XhoI and NheI and ligated with an insert containing a minimal promoter and a short stuffer sequence consisting of the first 100 bp of luciferase. The luciferase sequence is not functional and merely provides a transcript sequence linked to each oligo barcode, which is necessary for downstream sequencing library construction. The ligated plasmids were used to transform Stellar competent cells as described above. The final plasmid library pool was sequenced on an Illumina MiSeq to ensure an oligo library coverage greater than 90%.

#### Cell culture

Lentivirus was produced in 293T cells in 10cm plates. Cells were transfected with 3.75ug pUC MDG, 7.5ug pCMV Δ8.91, and 7.5ug plasmid library using Lipofectamine 2000 (Life Technologies). Viral particles were collected 48 hours post-transfection and concentrated using Lenti-X Concentrator (Takara). Lentivirus was titrated in primary keratinocytes to maximize viral transduction while minimizing lentiviral toxicity. For each MPRA biological replicate, 12 million keratinocytes were transduced in 15cm plates containing 5ug/mL polybrene. Cells were selected in 0.8ug/mL puromycin 24 hours post transduction. Once selected, cells were seeded for day 0, 3, and 6 timepoints of differentiation. At each timepoint, total RNA was isolated using Qiagen’s RNeasy Plus kit and then used to generate MPRA sequencing libraries.

#### MPRA sequencing library construction

cDNA was synthesized from total RNA using SuperScript IV (ThermoFisher Scientific) using a gene specific primer that anneals to the MPRA transcript. The primer also contains a 15 nt degenerate sequence that serves as a transcript UMI. cDNA synthesis reactions were cleaned up using SPRIselect beads (Beckman) and amplified using PrimeSTAR Max DNA Polymerase (Takara) for five PCR cycles to add Illumina sequencing adapters. Sequencing indexes were added in a second PCR step, which was monitored on a Stratagene MX3005P quantitative PCR machine to avoid library over-amplification. Final sequencing libraries were gel purified in a 2% agarose gel. Library concentration was determined using a KAPA Library Quantification Kit (Roche). Deep sequencing was performed on an Illumina NovaSeq 6000.

#### MPRA analysis

The DNA plasmid library was sequenced to capture the baseline fractions of each sequence in the library. Since the UMI is on read 1 and the barcode is on read 2 based on the primer locations, we perform paired ended sequencing. The reads are then trimmed (only the 20bps after the first 17 bps in read 2 constitute the barcode) and the UMI is associated with the read such that downstream analysis can proceed as single-ended data. These adjusted reads are aligned to the barcode sequences using bwa aln/samse with default parameters, and the aligned reads are then reduced by UMI to get unique read counts per barcode. These counts are then divided by the total to get the fractional value for each barcode in the library.

The MPRA library reads were sequenced and analyzed in the same fashion as the DNA plasmid library. The counts were then renormalized using the plasmid fractions by multiplying the MPRA counts by the plasmid fractions, converting to fractions, and multiplying by the total count across the MPRA library. In other words, the renormalization provides the counts per MPRA barcode assuming a uniform distribution of barcodes in the library at the same sequencing depth as was actually performed. These counts are then run through regularized log transform from DESeq2 to get a normalized signal matrix. This normalized matrix is then used in downstream analyses.

To test trajectory patterns, the normalized MPRA signal for all sequences belonging to the pattern are collected for days 0, 3 and 6. Then day 3 and 6 read outs are compared to day 0 by a Wilcoxon signed rank test (*p* < 0.05) to determine differential signal between timepoints. If the measurements show differential signal for any of the two days, the trajectory is considered to have dynamic activity across the time course. Then, the mean (across all sequences) pattern of the MPRA signal across the three time points is compared to the corresponding average ATAC trajectory to determine a correlative match (Spearman rank correlation *p* < 0.05) in terms of the dynamics.

To estimate interaction scores for motif pairs tested in the MPRAs, we compare the distribution of normalized MPRA signal (log scale) of endogenous sequences contained both motifs to the expected log-additive effect of each individual motif. When motif a is scrambled, we note the MPRA signal = a. When motif b is scrambled, we note the MPRA signal = b. When both motif a and b are scrambled, we note the MPRA signal =n. Then, the expected log-additive signal for the endogenous sequence containing both motifs = (a – n) + (b – n). We then utilize the Wilcoxon signed rank test (*p* < 0.10) to determine whether there is a significant difference between the observed endogenous signal and the log-additive expected signal. A significantly positive score indicates a super-multiplicative effect of the motif pair. A non-significant score indicates a multiplicative (log-additive) effect of the motif pair. A significant negative score indicates a sub-multiplicative effect of the motif pair.

### Analysis of genetic variation and heritability

We utilized LD-score regression software (https://github.com/bulik/ldsc) to determine genome-wide significant variants. From UK Biobank (http://www.nealelab.is/uk-biobank/), we utilized the GWAs results for the following phenotypes (codes in parentheses): basal cell carcinoma (20001_1061), eczema (200002_1452), psoriasis (20002_1453, L12_PSORIASIS, L12_PSORI_NAS, L40), non-melanoma malignant neoplasms of skin (C3_OTHER_SKIN, C44, C_OTHER_SKIN), actinic keratosis (L12_ACTINKERA), rosacea (L12_ROSACEA, L71), seborrheic keratosis (L82), diseases of skin and subcutaneous tissue (XII_SKIN_SUBCUTAN), and other/unspecified disorders of skin and subcutaneous tissue (L12_SKINSUCUTISNAS). We additionally utilized two other GWAS studies for dermatitis (GWAS catalog: GCST003184) and acne (GWAS catalog: GCST006640) as cited in the main text.

